# Interactions in self-assembled microbial communities saturate with diversity

**DOI:** 10.1101/347948

**Authors:** Xiaoqian Yu, Martin F. Polz, Eric J. Alm

## Abstract

How the diversity of organisms competing for or sharing resources influences community production is an important question in ecology but has rarely been explored in natural microbial communities. These generally contain large numbers of species making it difficult to disentangle how the effects of different interactions scale with diversity. Here, we show that changing diversity affects measures of community function in relatively simple communities but that increasing richness beyond a threshold has little detectable effect. We generated self-assembled communities with a wide range of diversity by growth of cells from serially diluted seawater on brown algal leachate. We subsequently isolated the most abundant taxa from these communities via dilution-to-extinction in order to compare productivity functions of the entire community to those of individual taxa. To parse the effect of different types of organismal interactions, we developed relative total function (RTF) as an index for positive or negative effects of diversity on community function. Our analysis identified three overall regimes with increasing diversity. At low richness (<12 taxa), potential positive and negative effects of interactions are both weak, while at moderate richness (12-20 taxa), community resource uptake increases but the carbon use efficiency decreases. Finally, beyond 20 taxa, there was no net change in community function indicating a saturation of potential interactions. These data suggest that although more diverse communities had overall greater access to resources, individual taxa within these communities had lower resource availability and reduced carbon use efficiency, indicating that competition due to niche overlap increases with diversity but that these interactions saturate at a specific threshold.

## Introduction

Organismal diversity is recognized as a driver of ecological functions such as biomass production, resource turnover, and community stability. As the number of taxa increases so do positive and negative biotic interactions such as facilitation, niche complementation, and competition, which all modulate the efficiency of resource use. While niche complementation leads to optimization of resource use through avoidance of resource use overlap, competition – either directly for resources or indirectly by chemical interference – often negatively impacts ecosystem functions. A considerable number of models have been developed to determine how and why the relative strength of different interactions on community function changes with diversity (Fox, 2006; Jaillard et al.; Jousset et al., 2011a; Loreau and Hector, 2001; Maynard et al., 2017). Most models determine the net effect of interactions on communities by comparing the observed community function to that predicted from monoculture functions of community members and agree on a general increase of all types of interactions with diversity, as well as diverse communities being more productive in many ecosystems due to the strong effect of niche complementation. It is, however, also possible for this relationship to be reversed (Cardinale et al., 2006). Especially in microbial systems, it has been proposed that the negative effects of antagonism on community function are only outweighed by the positive effects of niche complementation if the microbes are functionally dissimilar and the resource environment is heterogeneous (Jousset et al., 2011a). Recent work further shows that even when both conditions are satisfied, a negative relationship between diversity and biomass production may occur if interspecific competition is strong and hieratical, such as in a highly antagonistic system of wood degrading fungi (Maynard et al., 2017).

Because of their strong dependence on obtaining measurements of relevant functions of community members in monoculture, many community interaction models are most suitable for systems in which diversity can be varied by making a series of assemblages from well-characterized species. However, this approach is difficult for microbial ecosystems because they typically display high richness and often only a small portion of the total diversity can easily be isolated and grown in pure culture (Epstein, 2013). As a result, artificial microbial assemblages have been used to experimentally study organismic interactions and these assemblages have been limited in richness and phylogenetic diversity. Importantly, it is not clear to what extent such artificial assemblages reflect naturally occurring interactions, which are often the result of long term evolutionary processes of co-occurring organisms. It therefore remains an open question how different types of biotic interactions contribute to community function across ecologically relevant ranges of diversity for microbes that have co-diversified in the wild.

Here, we address the problem of how microbial interactions and their effects on ecosystem functions change over a wide diversity spectrum by generating dilution series of planktonic microbial communities and allowing them to self-assemble on seaweed extract as a realistic, complex environmental carbon substrate, which mimics an algal bloom in the coastal ocean (Takemura et al., 2017; Teeling et al., 2012). This process generated a series of replicate microbial communities with varying diversity and allowed measurement of biomass production and respiration as relevant community wide parameters. Following the measurements, each community was diluted to extinction to approach monoculture-level diversity, and the same production measures were obtained and compared to the community values. We extend relative yield total (RYT), a classical criterion for determining whether intercropping leads to higher crop yield than mono-cropping (De Wit and Van den Bergh, 1965), to our multi-taxa system and generalize it to summable functions across community members as the relative total function (RTF). RTF can be further broken down to investigate how interactions affect community production through changes in total resource use or resource conversion efficiency. We find that with increasing diversity, total resource access increases due to niche complementation, but carbon use efficiency decreases due to competition, leading to a stronger increase in community respiration than biomass production. Our results show that the production of a natural community may be limited by both its potential to access resources and by the amount of niche overlap that allows members to coexist in nature.

## Results

### Experimental Workflow

To test the relationship between diversity and community function, we generated microbial communities of decreasing diversity by serially diluting the same seawater sample and subsequently tracking relevant measures of microbial functions during growth in seaweed-seawater medium (SSM, pasteurized seawater containing extract from the brown algae *Fucus*). Specifically, our approach consisted of two stages where the first generated communities of similar overall biomass but different richness for which community production and respiration were measured, and the second consisted of dilution to extinction of communities from the previous stage to generate monocultures for which the same community functions were measured as input to the RTF model (Fig. 1, Methods).

**Figure 1.**
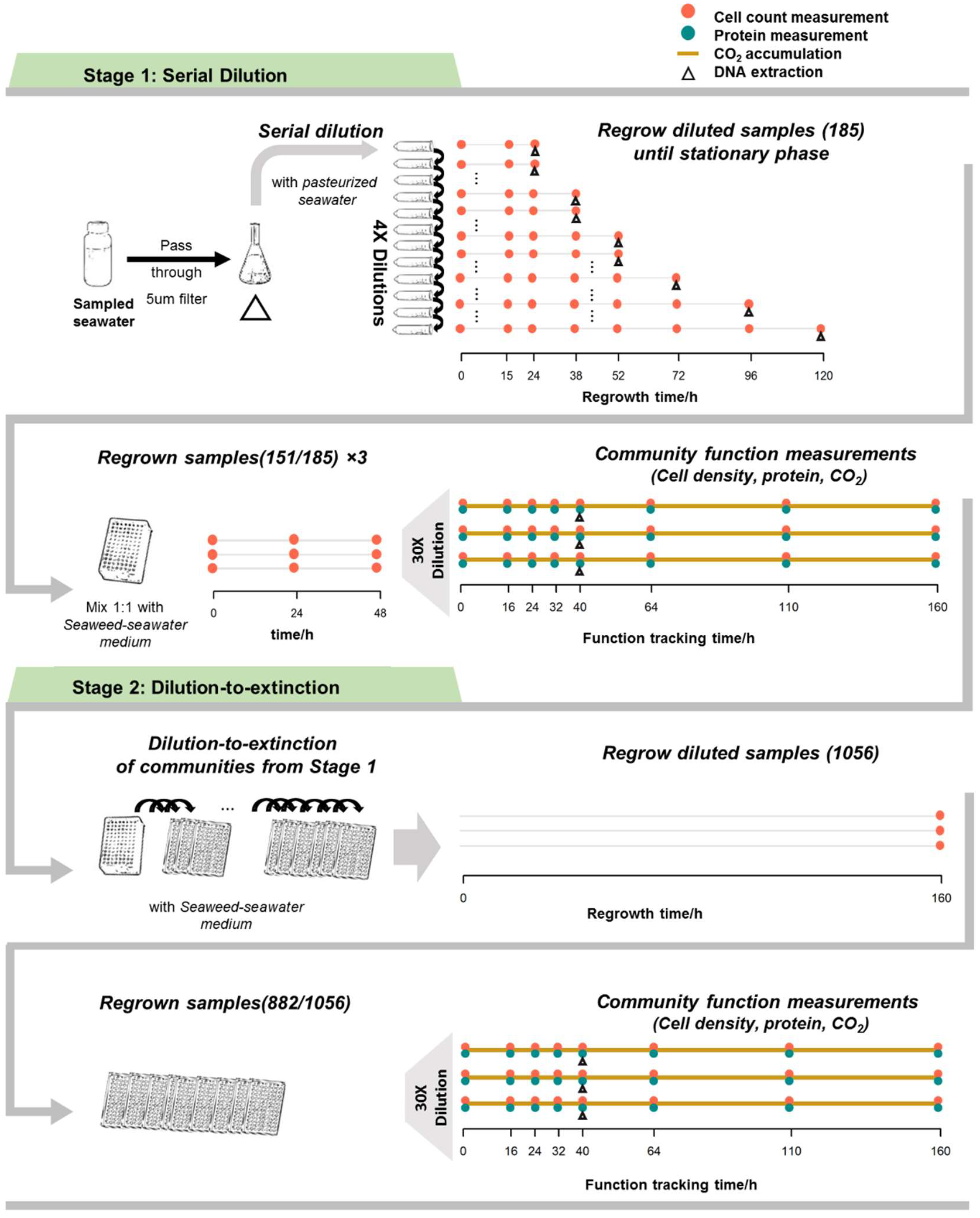
Schematic for community self-assembly and functioning measurements. In the first stage, a seawater sample is serially diluted and regrown in filtered seawater to generate inocula of different diversities for community growth and function measurements in seaweed-seawater medium (SSM) over a period of 160 hours. In the second stage, communities from the first stage are diluted to extinction to generate near monoculture samples that are subjected to the same growth and function measurements in SSM over a period of 160 hours.

In the first stage, we generated 15 replicate samples from seawater passed through a 5 μm filter and serially diluted each sample in 12 steps of 4-fold dilution each to yield a total of 185 communities of differing cell numbers and diversity. To acclimate these communities to the conditions used for measurements of community function, each was regrown first in pasteurized filtered seawater without any additional nutrients, then diluted 1:1 into SSM and regrown for 48h, and finally diluted 1:30 into SSM and grown for 160h (Fig. 1). This final regrowth experiment was used to measure community functions at different diversity levels. In the second stage, the resultant communities were diluted to extinction to obtain low complexity communities (typically 1-10 taxa) and regrown in SSM for 160h in order to determine the same functions under near monoculture conditions.

Biomass production was measured as the maximum of both cell numbers and per cell protein content that the communities reached within the 160h of observation, while respiration was estimated as the total CO_2_ production during this time. The combined measures of biomass production and respiration allow estimation of resource use efficiency. Finally, community composition was assessed by 16S rRNA amplicon sequencing immediately after the first diversity removal step and after each community reached early stationary phase in SSM. To estimate the total taxonomic richness, we first counted the number of taxa as clusters of identical 16S rRNA amplicon sequence variants (ASVs) (Callahan et al., 2016, 2017), and then inferred the number of ASVs that were uncounted due to limitations in sequencing depth (Willis and Bunge, 2015). Since observed and estimated taxonomic richness was highly similar in all cases (Fig. S1), we estimate richness from direct ASV counts.

Of the total 185 possible communities from the first stage dilution series, 151 showed sufficient cell numbers after regrowth in pasteurized seawater to serve as inoculum into SSM. These communities ranged in taxonomic richness and cell densities from 5 to 350 and 1.1×10^4^ to 3×10^5^ cells/mL, respectively (Fig. 1 and S2a). As expected, more diluted communities contained fewer and more dissimilar sets of taxa (Fig. S2b, c). All communities experienced further loss of taxa during their regrowth in SSM (Fig. S3a, b) possibly due to environmental filtering, population bottlenecks during transfer or competitive exclusion.

The second stage dilution to extinction experiment to create monocultures yielded 882 samples, most containing less than 10 taxa, with 275 almost completely dominated by a single taxon. These monocultures represent 37 ASVs in 7 families (Table S1), and cover 75.8% of total reads for the communities grown in SSM from the first dilution, hence providing a good basis for comparison of community-level and single taxon measures of function.

### Relationships between diversity and community functions vary

We observed Michaelis-Menten-like hyperbolic relationships between taxonomic richness and cell density or CO_2_ accumulation, but an inversely proportional relationship between taxonomic richness and protein production per cell, independent of whether taxonomic richness of the inoculum or resultant communities after growth in SSM was used (Fig. 2a, b). Normalizing both hyperbolic curves for respiration (CO_2_ production) and cell density to their maximum values, we found that respiration increased at a faster rate than cell density with increasing taxonomic richness (Fig. 2c, p=0.04, one-tailed t-test). Since the amount of protein per cell also decreased with diversity, this differential rate indicates that an increasing portion of assimilated carbon was released as CO_2_ as the number of taxa rises. Because such lowered yield in more complex communities could be due to individuals having intrinsically lower efficiencies in converting assimilated carbon into biomass or due to different taxa negatively affecting each other, we developed an indicator that allows differentiating between these two possibilities.

**Figure 2.**
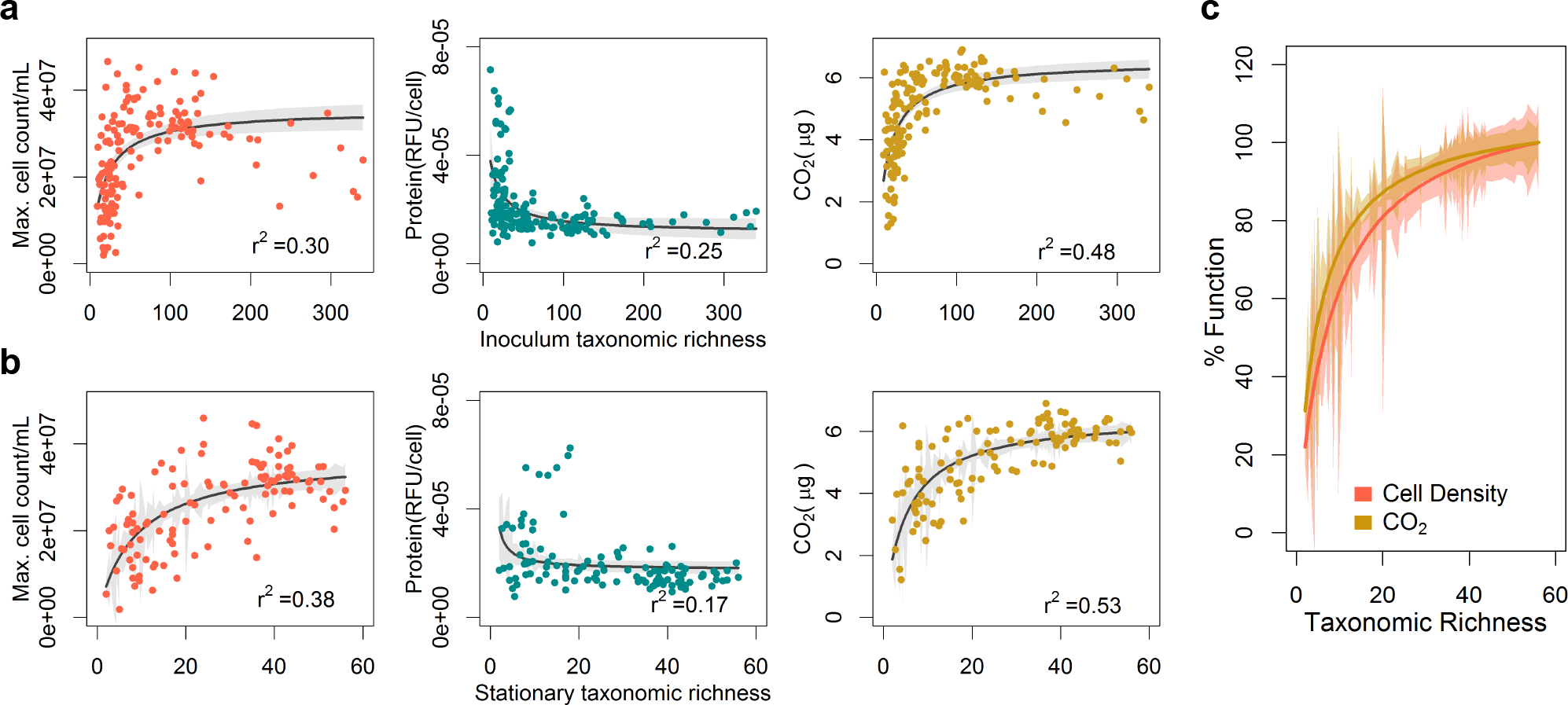
Diversity strongly impacts all measures of community function. Relationship of different measurements of community production (cell numbers and protein per cell) and respiration (CO_2_ production) with (a) inoculum taxonomic richness and (b) stationary phase taxonomic richness of communities. (c) Comparison of the hyperbolic fits for CO_2_ production and cell density scaled to 1 as maximum. Each dot represents one inoculum community; blac/colored lines indicate fits to an appropriate model with the least AIC between a linear, log-linear, and hyperbolic least squares fit, with gray/colored regions around the line indicating the 95% confidence of the fit.

### Relative total function as an indicator for interaction effects on community function

We established relative total function (RTF) as an indicator for inter-taxa relationship effects on community function by generalizing the concept of relative yield total (RYT) suggested by De Wit & Van den Bergh (1965). RYT is calculated as the sum of the relative yields of two species in a community compared to each of their monocultures. It is thus a measure of how resource use of one species is influenced by the other when the two occur together in a community, under the assumption that the resource to biomass conversion efficiency of each species is maintained. Generalizing two species to multi-species and yield to all community functions that are summable across species, we defined RTF as the summed relative function of each taxon in a community to their monocultures:

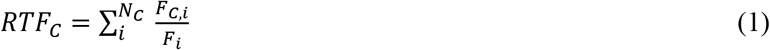

where *F*_*C,i*_ is the function of a taxon *i* in community *C*, *F*_*i*_ is the measured monoculture function of taxon *i*, *N*_*C*_ is the number of taxa in community *C*. *F*_*C,i*_ is calculated by multiplying the measured total community function *F*_*C*_ with relative abundances of each taxon from community composition measurements (see Methods for related assumptions). Under the null model that community function is solely dependent on individual function, *RTF*_*C*_ equals 1, while a *RTF*_*C*_ larger than 1 indicates that biotic interactions increase community function, and a *RTF*_C_ smaller than 1 indicates a decrease (see Fig. S4 for graphical illustration).

Under most resource concentration regimes, the carrying capacity of a community can be seen as a linear function of resource uptake. Only when resource overabundance induces incomplete respiration of substrates, the assumption of linearity may be violated (Polz and Cordero, 2016). Since this is unlikely here due to limited resources being provided, we can further define for community functions analogous to carrying capacity:

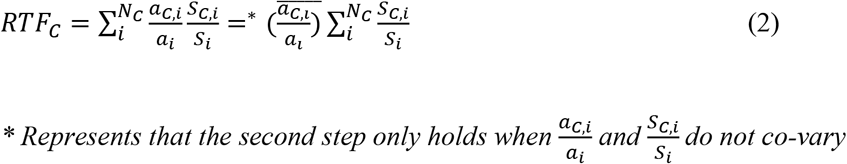

where *a* is efficiency of resource conversion into the function of interest and *S* is the amount of resource uptake, meaning there are two major ways that interactions could affect community function: by altering the total amount of resource uptake or the efficiency of resource conversion into the community function of interest. Under the null model, both the mean relative resource conversion efficiency 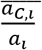, and the relative total resource uptake 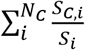 would be 1. If either factor is larger or smaller than 1, it is positively or negatively affected by interactions between taxa.

While *RTF*_*C*_ compares the function of a community to the monoculture functions of its constituents, *RTF*_*C*_/*N*_*c*_ compares the function of individual taxa in a community to its monoculture.

A *RTF*_*C*_/*N*_*c*_ larger than 1 is a signal for facilitation between the majority of community members. Since *RTF*_*C*_ is the product of relative total resource uptake and relative resource conversion efficiency, *RTF*_*C*_/*N*_*c*_ can be seen as the product of mean relative per taxa resource uptake 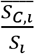 (which in ecological terms, is the ratio of the realized niche to the fundamental niche), and mean relative resource conversion efficiency 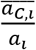.

We used the *RTF*_*C*_ indicator to evaluate how interactions between taxa impacted different community function measurements as diversity increased. Since *RTF*_*C*_ is only suitable for community functions that are summed across taxa, we applied it to three functions of interest: cell number, total protein production, and total respiration (as CO_2_ accumulation). However, calculation of *RTF*_*C*_ also requires knowing the monoculture function of every taxon in the community. We achieved this by defining communities as “constitutable” that were at least to 85% covered by the 37 taxa we were able to isolate in sufficient purity. Hence we assumed that the remaining 15% of taxa contribute to community function similar to the constitutable 85%. The 85% cutoff was chosen because it was the highest criterion that allowed constitutable communities to cover the full diversity spectrum of the stationary phase communities (Fig. S5). Using this cutoff, we identified 82 (35%) of the 235 total communities with good sequencing coverage to be constitutable for every community function.

### Interaction effects on community function differentially increase with diversity

By tracking how different community functions relative to monoculture change with diversity, we found that as diversity increased, interactions lead to a stronger increase in community respiration (CO_2_ production) than community biomass production (cell count and total protein). At low richness (*N*_*c*_ ≤ 12), *RTF*_*C*_ was not significantly different from 1 for CO_2_ production, and slightly below 1 for total protein production and cell count, indicating that the effect of interactions on respiration was negligible but weakly negative for biomass production (Fig. 3a, S6a, t-test, two-tailed, p_CO2_=0.95, p_protein_=1.04×10^−6^, p_cell_=2.43×10^−10^). However, as taxonomic richness increased beyond 12, *RTF*_*C*(*CO2*)_ steadily rose above 1 until it eventually plateaued when a moderate richness level of 20 taxa was reached, while *RTF*_*C*(*Protien*)_ remained around 1 over the entire diversity range, and *RTF*_*C*(*Cell*)_ remained around 1 until 20 taxa, beyond which it stayed slightly above 1 (t-test, two-tailed, p_CO2,12<Nc≤20_=0.02, p_CO2, Nc>20_=5.09×10^−11^, p_protien,12<Nc≤20_=0.81, p_protien, Nc>20_=0.35, p_cell,12<Nc≤20_=0.24, p_cell,Nc>20_=0.06). Thus, at moderately high diversity, interactions had a strong net positive effect on community respiration, but only a weakly positive effect on the production of cells, and no net effect on community protein production.

**Figure 3.**
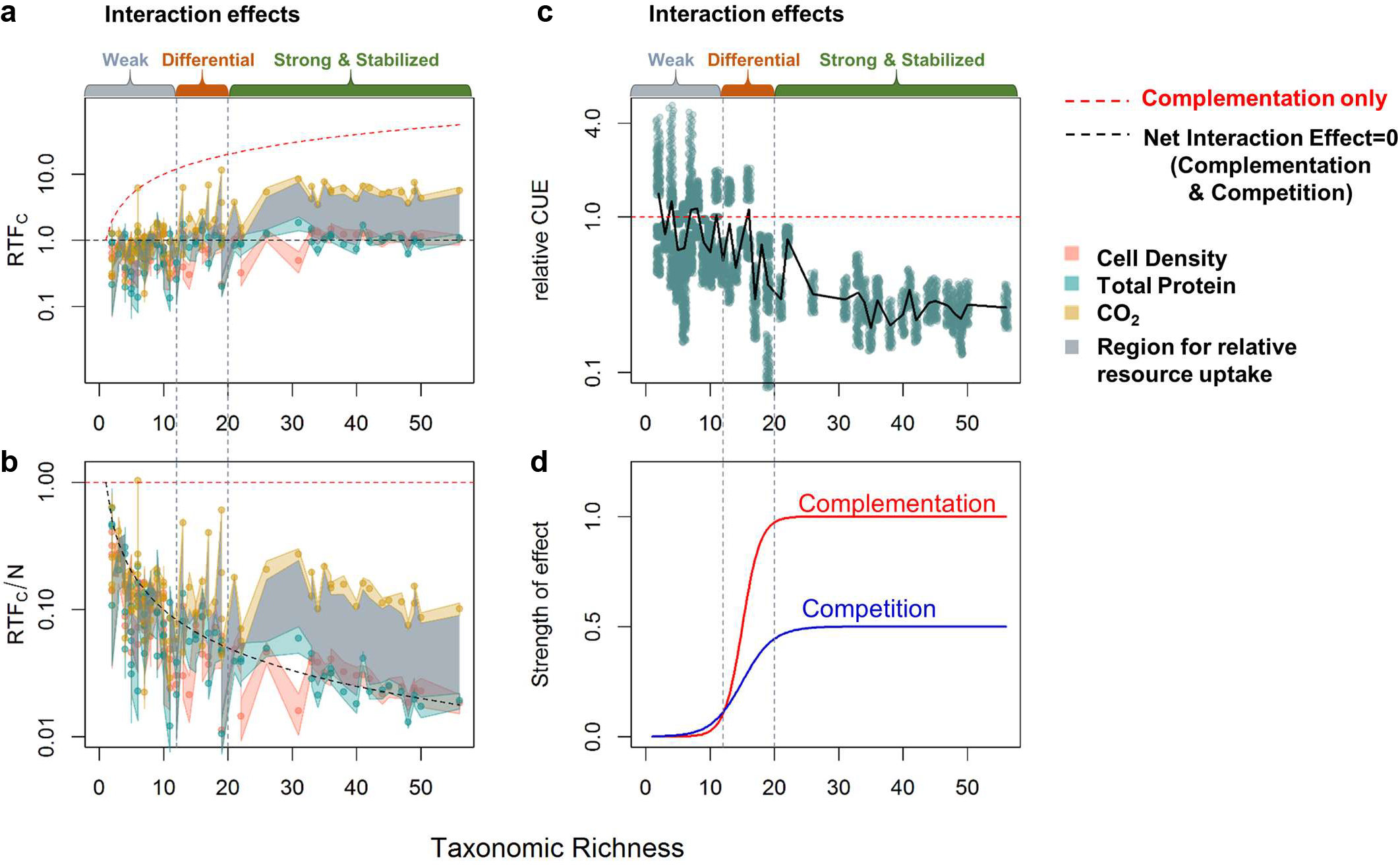
Relative community-to-monoculture functions indicate differential increases in niche complementation and competition with diversity. Relationship between diversity and (a) relative total biomass production, respiration, and resource uptae, and (b) relative per taxa biomass production, respiration, and resource uptae (c) estimated relative carbon use efficiency (CUE). In (a), (b) each point represents one community at stationary phase (biological replicates were not averaged). In (c), each point represents a combination of one relative total function (*RTF*_*C*_)ratio with a random value of CUE drawn from 0-0.6. The blac line represents the mean of all combinations with the same taxonomic richness. Colored regions around the points indicate the standard deviation of the interaction effect, and the grey areas indicate the range where relative total and per taxa resource uptae is limited to at each taxonomic richness. The y axes are in log scale for better resolution of points (see Fig. S6 for the same data in linear scale). (d) A hypothetical model for how the effects of niche complementation and competition on community function scale with taxonomic richness. Numbers on the y-axis are arbitrary.

### Community resource uptake increases with diversity while individual taxa resource uptake decreases

Since the relative community CO_2_ production increased significantly with diversity, and relative community biomass production also exhibited a weak increase, the relative resource uptake of the community must also increase with diversity. Detailing this under the *RTF*_*C*_ framework, since each community has a single relative total resource uptake value 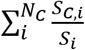, and the conversion efficiencies for CO_2_ and biomass accumulation have to change in opposite directions (i.e., they cannot simultaneously increase or decrease), the relative total resource uptake of a community must fall between *RTF*_*C*(*CO2*)_ and *RTF*_*C*(*Biomass*)_. Taking total protein production as the measurement for biomass, we find that the region between *RTF*_*C*(*CO2*)_ and *RTF*_*C*(*Protien*)_ move from around 1 to higher than 1 as diversity increases (Fig. 3a, S6a). This indicates that biotic interactions with positive effects on resource uptake, such as niche complementation, are more prevalent in more diverse communities.

However, individual taxa in more diverse communities on average take in fewer resources, i.e., they have narrower realized niches relative to their fundamental niches in monoculture. This is supported by the gradual decrease of 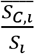, whose boundaries are defined by *RTF*_*C*_/*N*_*c*_ and *RTF*_*C*(*Protien*)_/*N*_*c*_, with diversity (Fig. 3b, S6b). Also, since all *RTF*_*C*_/*N*_*c*_ never exceeded 1, communities where most community members facilitate each other are rare. In fact, only a very small fraction of taxa produced more biomass or CO_2_ in communities than in monocultures (Figure S7), indicating that each taxon always has more competitors than facilitators, and the frequency of mutualistic interactions among different taxa is lower than negative interactions.

### Competition in more diverse communities decrease carbon use efficiency (CUE)

Since relative community CO_2_ production increased faster with diversity than relative community biomass production, the relative carbon to biomass conversion efficiency (often known as carbon use efficiency, CUE) decreased, possibly as a result of stronger competition. Since the ratio of *RTF*_*C*(*CO2*)_ to *RTF*_*C*(*Biomass*)_ should scale positively with both the expected CUE from monocultures and how much this CUE changes upon introduction of the organism into a community, we estimated the range of the relative-to-expected CUE from the *RTF*_*C*(*CO2*)_ to *RTF*_*C*(*Protien*)_ ratios and limiting the expected CUE to the thermodynamic limits of 0-0.6 (Geyer et al., 2016, see Methods for details). We found that as taxonomic richness increases, interactions had increasingly negative effects on the relative CUE of the community, but the effect leveled off at moderate taxonomic diversity (Fig. 3c, S6c). Furthermore, since at higher richness *RTF*_*C*(*Protien*)_ was lower than *RTF*_*C*(*Cell*)_ (*N*_*c*_ > 20, p=0.08, Paired Wilcoxon rank sum test, one-tailed), the relative CUE decrease is a joint effect of both smaller cell size and lower cell number. This is likely because microbes in more diverse communities are under stronger competition for resources, as evidenced by their narrower relative realized niches. They either have to grow faster to directly compete with other microbes for preferred resources or avoid competition by using more recalcitrant and less preferred resources. Because growth rate generally scales positively with cell size (Cermak et al., 2017; Roller et al., 2016; Schaechter et al., 1958), it appears more likely that our observation of smaller cells in more diverse communities indicates reduced growth rate and partitioning of a larger portion of carbon taken up toward respiration due to the lower energy yield of more recalcitrant substrates.

However, it is possible that the “faster growth” strategy dominates initial periods of growth and the “recalcitrant resource” strategy takes over in later growth phases. We observed that more diverse communities had, at least initially, faster growth rates. An overwhelmingly majority of communities with more than 20 taxa reached maximum cell density by 24h of growth, while those with less than 20 taxa had equal probability of growing to maximum at 24h, 32h or 40h (Fig. 4). Hence, our data suggest that diverse communities had a high probability of containing members specialized for substrates that allow initial rapid growth. Conversely, communities with lower diversity only occasionally contained such potentially fast growers, explaining the broader distribution of times to maximum cell numbers. Although this does not provide direct evidence for individual taxa growing faster in more diverse communities, it may indicate that it could be advantageous for taxa in more diverse communities to regulate itself for faster growth compared to monocultures. We would then expect relatively lower CUE but larger cells to accumulate early in the experiment (Lipson, 2015; Pfeiffer et al., 2001). Over the full observation period, however, more diverse communities on average had smaller cell size, possibly because after reaching peak density, the initial population is replaced by other taxa or switched into a metabolism that utilizes substrates that are more recalcitrant and less energy efficient.

**Figure 4.**
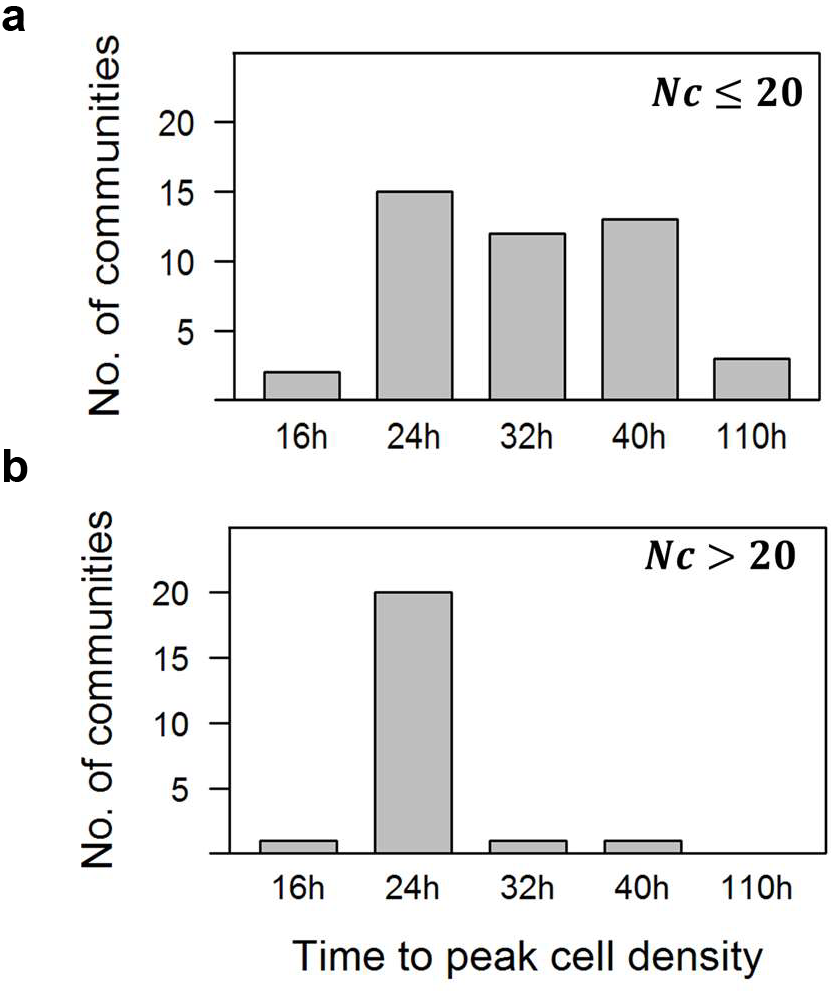
More diverse communities reach stationary phase earlier. Distributions of time for communities to reach pea cell density in communities with (a) less than 20 taxa, and (b) more than 20 taxa.

### Effects of interactions on community function increase logistically with diversity

Although cell density and CO_2_ both exhibit Michaelis-Menten-like hyperbolic relationships with diversity, both competition and complementation have logistically increasing effects on community function as diversity increases. By estimating CUE through the size of the difference between *RTF*_*C*(*CO2*)_ and *RTF*_*C*(*rProtien*)_ and total niche occupation through their relative position to 1, we found that at low diversity (*N*_*c*_ ≤ 12), the effects of both competition and niche complementation were weak, with the negative effects of competition narrowly exceeding benefits of niche complementation. As diversity increased beyond this level, the positive effect of niche space expansion gradually exceeded that of the negative effect of competition until they both stabilized at moderately high diversity (*N*_*c*_ > 20). This indicates that while niche complementation is probably limited by the total amount of resources available, competition also has plateaued (Fig. 3d). Overall, these patterns translate to a logistic model of growth as diversity increases for impacts on community function by either competition or complementation.

### Specific taxa effects on community function

Since it is often assumed that organisms that are more phylogenetically distant are also more metabolically distinct, and since taxonomic richness only partially explained the variance of community function (Fig. 2a), we asked if phylogenetic diversity of the communities affected community cell production or respiration through niche complementation. We found that neither of the two most common measurements for phylogenetic distance, the abundance weighted mean pairwise distance (MPD) or the abundance weighted mean nearest taxon distance (MNTD), had significant impact on community function, by performing two-way ANOVA with the logarithm of taxonomic richness and MPD or MNTD as factors (MPD: p_Cell_ =0.04, R_cell_^2^=0.02; p_CO2_=0.54; MNTD: p_cell_=0.14; p_CO2_=0.004, R_CO2_^2^=0.04).

We then checked the possibility of specific “key” taxa affecting community function, i.e., whether one taxon could alter community function without altering MPD or MNTD. We screened for these taxa by looking for ASVs whose relative abundance significantly correlated (Kendall rank correlation, q<0.05 after FDR correction) with community function within sliding windows of taxonomic richness (window widths ranging from 2-10) (see Table S2 for all significant correlations). In most cases, each identified taxon was specific to a certain range of taxonomic richness (usually of size 10-15), with the direction of correlation highly consistent among windows within the range. Also, correlations between taxa and CO_2_ were few compared to those for cell density and total protein production, indicating that biomass production is more sensitive to competitors than respiration. We identified five ASVs that exemplify how diversity may affect the outcome of microbial interactions and consequently community function.

Taxonomic richness of the inoculum strongly influenced dominance patterns in communities and thereby had significant effects on community function. For example, one taxon (ASV3) belonging to the genus *Alteromonas* was found to positively correlate with CO_2_ production but only in communities with less than 10 taxa. In most of these communities, ASV3 either completely dominated or was close to 0 in abundance, indicating that ASV3 is very likely a fast-growing generalist capable of utilizing a wide range of substrates and thereby outcompeting other microbes. ASV3 could be similar to the “super-generalist” *Alteromonas* strain AltSIO, which was found to be capable of consuming the entire pool of labile DOC in coastal surface seawater (Pedler et al., 2014). However, ASV3 did not show strong dominance in communities with higher inoculum diversity. This suggests that there were other, more rare taxa that were able to compete with ASV3 and demonstrates the importance of priority effects in colonization of small resource patches in the environment (Datta et al., 2016).

We also observed that one taxon could have opposite correlations with community function at different levels of community diversity. As richness increased beyond ~20 taxa, ASV18 (a type of *Polaribacter*) switched from being positively to being negatively correlated with cell density. Since some *Polaribacter* species are broad spectrum degraders of algal polysaccharides (Xing et al., 2015), the switch in correlation may indicate that at low richness ASV18 might have occupied a niche with few competitors, or even played a facilitative role by degrading a complex polysaccharide. This process requires secretion of polysaccharide lyases and may induce cross-feeding in the community due to incomplete uptake of the hydrolysis products by the primary degrader (Datta et al., 2016; Hehemann et al., 2016). At higher richness, however, increased competition for the same resource may have led to more inefficient growth.

Despite having generally more niche overlap for resources, communities with high diversity can still benefit from having taxa that use the less common resources. ASV30, identified as *Wenyingzhuangia*, positively correlated with total protein production in communities with high taxonomic richness. Members in the *Wenyingzhuangia* genus have been identified as specialists for degrading sulfated polysaccharides (Shen et al., 2017) and are one of the few organisms known to degrade the highly sulfated and recalcitrant sugar Fucoidan (Chen et al., 2016), estimated to be 4-10% of the dry weight of *Fucus* (Fletcher et al., 2017). ASV30 may thus have positively affected community productivity by occupying a niche inaccessible to most other taxa, and might even have acted as a pioneer taxon by degrading the recalcitrant Fucoidan and converting it into more easily usable substrates.

Other taxa were found to have effects on community production likely through facilitation and predation. Several ASVs belonging to the genus *Sulfitobacter* were consistently found to be positively correlated with community productivity across different ranges of community richness and productivity measurements. Members of this genus are found as stable associates with algae (Singh and Reddy, 2014), and possess sulfite-oxidation and aromatic compound degradation abilities (Mas-Lladó et al., 2014). Given that *Fucus* contains large amounts of aromatic compounds, *Sulfitobacter* could have positively affected the community productivity through degradation of aromatic compounds, which could otherwise impede the growth of other bacteria. We also found a negative correlation between ASV66, a putative predatory bacterium from the genus *Halobacteriovorax*, and community cell density at high taxonomic richness, indicating that predatory behavior may also play a role in determining community production.

## Discussion

In this study, using self-assembled microbial communities directly derived from a costal ocean seawater sample, we examined how diversity could impact community function through various interactions. Consistent with observations from artificially assembled systems, we found communities with greater diversity to have greater resource uptake due to niche complementation. However, the expansion of resource use came at the cost of increased competition driven by niche overlap and decreased the carbon use efficiency of the communities.

Compared to assembling “bottom-up” communities from known isolates, our method has both advantages and caveats. Since our communities self-assemble, our workflow is inevitably less well controlled compared to artificial assembly experiments and has more challenges regarding detailed characterization of individual traits and interactions. For example, our inoculum communities had cell densities on the same order of magnitude, and although multi-variable regression showed that relationships between diversity and community function remained significant after considering the effect of initial cell density (Table S3), we were not able to control them to be the same exact cell density as done in isolate assemblages. Also, since our method was based on 16S rRNA gene amplicon sequencing, we were only able to distinguish bacteria that had at least one nucleotide difference in the 16S rRNA V4 region. Different strains of bacteria that show genetic variability beyond the 16S rRNA region and have different productivities would be collapsed as one taxon in our analyses.

Despite these limitations, our method has the advantage of providing a more accurate recapitulation of how natural microbial communities assemble under defined environmental conditions and a more precise measurement of how community diversity alters different productivity measurements. This is especially important in light of studies that show evolutionary history can alter resource use patterns of taxa and have a strong influence on how community functions are affected by biodiversity (Fiegna et al., 2014; Gravel et al., 2011). While artificially assembling communities from isolates allows exploration of theory behind the relationships between diversity and ecosystem functioning, they are less suitable for evaluating the effects of diversity on ecosystem functioning in nature. With isolation being an extreme population bottleneck, the evolutionary history of communities being assembled are completely different from that in natural communities. We thus argue that a “top-down” approach of directly taking natural communities to self-assemble to different diversities, and then parsing the behavior of the self-assembled communities provides a better window to look at how diversity affects community function in nature.

In our system, we found a large decrease in CUE with diversity, possibly due to increased competition for resources. Although decrease of carbon use efficiency (CUE) with increasing diversity due to interactions has been documented previously, our data suggest an unusually large effect. For example, in an artificially assembled system of saprophytic basidiomycete fungi, interactions in multispecies communities were found to decrease CUE by up to 25%, a stronger reduction than induced by many abiotic factors such as temperature increase (Maynard et al.). However, most of our communities at moderately high diversity were estimated to have a 60%-80% decrease in CUE due to interactions. While the CUE decrease in Maynard et al was largely attributed to antagonism, it is unlikely the case in our system, since instead of selecting a system (wood decaying fungi) and conditions that favor antagonism, we are mimicking a free-living and dilute marine environment where antagonism is likely less of a factor. Although antagonistic potential has been demonstrated in marine bacteria (Cordero et al., 2012; Rypien et al., 2010), it is more likely that the large interaction effect on CUE in our system results from resource competition, which Maynard et al made efforts to minimize by having excess supply of both carbon and nitrogen. With seaweed extract mostly being carbohydrates, our system is probably limited by nitrogen or phosphate; thus with the narrower realized niches we observe as diversity increases, resource competition is likely to be prevalent in our communities.

Since our communities originate from natural marine communities, the large niche overlaps they exhibit when self-assembling on SSM, as well as the saturation of community production at moderate taxonomic richness, are reflective of the ubiquity of algal exudates in the coastal environment. Where a positive but decelerating diversity-productivity curve starts to plateau is often seen as an indicator for substrate generality, i.e., less organisms will evolve and maintain pathways for the degradation of a substrate if it is rare, and thus more organisms will be needed to saturate the niche space that the substrate creates (Delgado-Baquerizo et al., 2016). However, the large niche overlaps also suggest that the concentration of algal exudates undergoes temporal fluctuation, since in an environment where substrates are at stable concentrations, organisms would evolve to differentiate niche occupation, as observed in assemblages of tree-hole bacteria grown continuously in tree leachate (“beech tea”) (Fiegna et al., 2014). Had our communities come from such an environment, adding more taxa might have increased the total resource uptake of the community but not increased competition for resources.

Furthermore, in our system, the impacts of both competition and complementation on community function logistically increased as diversity increased, probably as a result of algal exudates being a complex mixture of substrates that have different numbers of bacterial consumers in the coastal ocean. Certain substrates may only be utilized by a portion of bacteria in the environment, and the chance of getting such bacteria in a community would be equally small among communities with different richness when diversity is low (*N*_*c*_ ≤ 12). Only when there was a sufficient number of taxa in the community did adding in new taxa actually expand the resource use profile, and this expansion eventually become limited by the total amount of resources available. Moreover, the speed of the resource use expansion was slower than that of the increase in taxa, thus niche overlap also increased with diversity. However, niche overlap also only started to translate to negative effects on community production when there was a sufficient amount of taxa in the community, possibly because only when there are enough competitors around there would be a need for employing a fast growth, low efficiency strategy.

Competition in our system saturated at moderately high diversity, possibly due to a combined effect of organisms co-diversifying over long periods of time under the specific environmental conditions of the coastal ocean. The limited amount of competition we observe is consistent with theoretical predictions that environmental fluctuation place upper bounds on how much overlap there can be between niches for species in the community to stably coexist (May and MacArthur, 1972). Indeed, the communities assembled here were drawn from the highly fluctuating coastal environment, and given that niche overlap is the result of organisms having sets of redundant functional genes, this might indicate that there is a limited amount of functional similarity between algal degrading bacteria due to the frequency of environmental fluctuation. While the observations we have in our system are specific to the coastal marine environment, we believe that they imply that in many natural habitats, there are general links between the stability of the environment, the amount of functional redundancy that it could support, and community production.

## Acknowledgements

This work was supported by the U.S. Department of Energy (DE-SC0008743) to M.F.P. and E.J.A. X.Y was partially supported by a Lord Foundation Graduate Fellowship. We are grateful to Christopher Corzett for providing us dried *Fucus* and for helpful discussions, as well as the BioMicroCenter at MIT for their assistance with sequencing, and the Koch Institute Flow Cytometry Core for their assistance on flow cytometry. We also wish to thank Sean Gibbons, Fangqiong Ling, and Joseph Elsherbini for helpful discussions and/or comments on an early version of this manuscript.

## Methods

### Media preparation

To prepare pasteurized seawater as a media for bacterial growth and as the solvent for making seaweed-seawater media (SSM), a total of 8L of costal surface seawater was collected from a sampling site near Northeastern University’s Marine Science Center (Canoe Beach, Nahant, MA, USA; N 42° 25’ 11.6″,W 70° 54’ 24.8″), on Nov 12^th^, 2016. The water temperature at the time of sample collection was 12.0°C. Seawater was pasteurized as described in Takemura et al 2017. Briefly, the seawater sample was divided into 2L bottles, heated to temperatures between 78-82°C in water baths and maintained at the temperature for 1h. Each bottle was pasteurized twice with at least 48h intervals between the pasteurization events. The pasteurized seawater was then combined and filtered through 0.22um filters to remove any large size particles.

Stock solution for making the SSM was made from *Fucus vesiculosus* collected from the rocky shorelines of Canoe Beach on July 12^th^, 2015. The *Fucus* was washed, sun dried and grinded using a blender (Waring). Four grams of the ground *Fucus* was mixed with 100mL of pasteurized seawater, and stirred at 150rpm for 2h at room temperature. The mixture was then passed through 20um Steriflip filters (Millipore), diluted 4 fold with pasteurized seawater, passed through a 0.22um filter (Corning, prewashed three times with MilliQ water), and pasteurized again. The seaweed media extract stock solution thus was 1% (w/v) and was stored in the dark at room temperature till time of use, when it was diluted to 0.1% (w/v) in pasteurized seawater to make SSM.

### Sample collection and experimental design

In order to generate the inoculum communities, a costal surface seawater sample was collected from a sampling site near Northeastern University’s Marine Science Center (Nahant, MA, USA; N 42° 25’ 11.6″,W 70° 54’ 24.8″), on Nov 18^th^, 2016. The water temperature at the time of sample collection was 11.5°C.

The collected seawater was filtered through a 5um filter (Whatman) to remove particulates and larger eukaryotes and was estimated to have a microbial concentration of 3*10^5 cells/mL via Fluorescence-activated cell sorting (FACS) using absolute count beads. Thus, an initial “undiluted” seawater community was defined as 30mL of the filtrate, containing ~10^6 cells in total. Fifteen “undiluted” seawater communities were 4-fold serial diluted with pasteurized seawater, generating 15 communities (30mL) for each dilution level (1/4^2^ to 1/4^11^). An additional 5 and 15 sub-communities were generated for the two highest dilution levels (1/4^10^ and 1/4^11^).

All communities were placed in 50mL Falcon tubes and rotated end-over-end at 6.5 rotations per minute in the dark. Bacterial growth in the tubes were tracked over time using FACS, with time points taken at 0, 15, 24, 38, 52, 72, 96, 120 and 360h. Communities were sampled for DNA extraction and inoculation when determined to have reached stationary phase, i.e., less than 20% increase in cell count between two consecutive time points following time points with over 20% growth in between.

Cell densities in all sampled communities ranged between 1.1×10^4^ to 3×10^5^ cells/mL. For each community, three replicates of 0.3mL each were used to inoculate an equal volume of 0.2% seaweed extract media. The communities were allowed to grow for 48h in 96 deep well plates on a floor shaker at 300rpm before being diluted 1/30 into 580uL of 0.1% seaweed extract media. The re-inoculated cultures were grown in MicroResp Systems (James Hutton Ltd, Aberdeen, UK) on a floor shaker at 300rpm, and their community functions tracked as cell count, total protein, and CO_2_ production at time points 0, 16, 24, 32, 40, 64, 120, and 160h. The 160h data point for cell count and protein was eventually omitted due to possible changes in the physical properties of the culture interfering with the FACS measurement.

At the end of the tracking period, communities similar in initial dilution levels and 160h cell count were combined. The combined communities were diluted in 0.1% seaweed extract media according to their cell densities so that on average each diluted community would contain 1 cell/200uL of culture. Each combined community had 18-24 corresponding diluted communities. These diluted communities were allowed to grow for 7 days in flat-bottom 96 well plates before they were screened for positive growth using FACS.

Communities that scored positive for growth were diluted 1/30 into 580uL of 0.1% seaweed extract media, and again grown in MicroResp Systems on a floor shaker at 300rpm, with their community functions tracked as cell count, total protein, and CO_2_ production at time points 0, 16, 24, 32, 40, 64, 120, and 160h. The 160h data point for cell count and protein was eventually omitted due to possible changes in the physical properties of the culture interfering with the FACS measurement.

### Community growth and function measurements

During regrowth of the diluted communities in pasteurized seawater for generating the inoculum communities, for each time point, 100uL subsamples of the regrown communities were obtained and fixed 1:1 with 0.8% Formaldehyde+0.5ug/mL 4’,6-Diamidino-2-Phenylindole (DAPI, Sigma). Cell counts were determined using a BD LSR II Flow cytometer with an activation wavelength of 355nm and 450/50nm band-pass filter. Cells that were between 400-2*10^4^ FSC-H, 20-10^5^ SSC-H and showed more than 200U fluorescence were counted as bacteria. All bacterial counts were normalized against 6 standards that contained CountBright absolute count beads (Thermo Fisher) at 990,000 beads/mL.

Communities inoculated into SSM were co-measured for cell count and protein per cell according to the methods described in Zubkov et al 1999 with some modifications. Briefly, at each time point 20uL of each culture was fixed 1:10 with 0.8% Paraformaldehyde (BeanTown Chemical). The fixed cultures were mixed 1:1 with staining media (1:5000 SYPRO red+0.02% SDS+1ug/mL DAPI) for 30min in the dark at room temperature before being run through a BD LSRFortessa Flow Cytometer with a high throughput sampler. Cell counts were determined using an activation wavelength of 405nm and 450/50nm band-pass filter. Total protein was determined by obtaining average fluorescence per cell under a 561nm laser and 610/20 band-pass filter and multiplying the fluorescence by cell count. For each 96 well plate of samples, cell counts were normalized to 3 standards of CountBright absolute count beads (Thermo Fisher) at 990,000 beads/mL, and total fluorescence was normalized to 3-6 wells that contained a fixed standard marine bacteria mixture containing approximately 1:1 *Vibrio* and *Halomonas*.

CO_2_ production of communities was calculated from reading indicator plates in the MicroResp System on a plate reader at λ=572nm. In MicroResp system, all target communities were placed in deep 96 well blocks and hermetically connected to a top indicator plate using a seal. The seal insulated wells from each other but allowed the indicator plate to reflect the % of CO_2_ accumulated in the headspace of each well. The MicroResp indicator plates were made and calibrated according to the manufacturer’s instructions. The relationship (R^2^=0.996) between %CO2 in headspace and absorbance was (%CO2) =0.1648/ (Δ_572_-0.2457)-0.2301, with Δ_572_ being the difference in A_572_ between the start and end of CO_2_ production time.

The rate of CO_2_ production per volume of culture was calculated from %CO2 for each sampling interval, and a normalized total CO_2_ production of communities was calculated by summing CO_2_ production rate* average time*average community volume for each sampling interval. The effect of atmospheric CO_2_ was removed by normalizing the CO_2_ production values to the average of the blank wells.

Measurements eventually used as community functions were: maximum cell density the community reached within 160h, maximum total protein production and protein per cell within 160h, as well as total normalized CO_2_ production within 160h. To account for cells clumping in later time points, the maximum protein per cell was calculated from maximum total protein production/maximum cell density, instead of directly comparing protein per cell measurements between time points.

### DNA extraction

In order to determine community composition for the inoculum communities, 30mL of the inoculum communities were pushed through Swinnex Filter holders (13mm, Millipore) containing 13mm 0.22um filters (autoclaved, Durapore membrane PVDF, Millipore) connected to Luer-Lok syringes (BD). Filter paper was removed from the holder, cut into 4-6 smaller pieces, submerged in 125uL QE buffer with 1% Ready-Lyse Lysozyme (Epicentre, Quick Extract Kit) in eppendorf tubes, and shook at 400rpm overnight at room temperature. The tubes were spun down at 1700rpm for 5mins the second day and the supernatant was stored at −20°C till future use.

The composition for communities growing in SSM were determined at early stationary phase. For each community, 200uL of sample was taken and filtered through MultiScreen HTS GV filter plates (0.22um, sterile, PVDF membrane, Millipore) by spinning the plates for 5 mins at 3000rpm. Each well was incubated overnight in 100uL QE buffer with 1% Ready-Lyse Lysozyme (Epicentre) on a tabletop shaker at 400rpm. DNA extract was collected by spinning the plates for 5mins at 3000rpm and obtaining the flow through.

### Library Prep and Sequencing

16S rRNA gene amplicon libraries (V4 hypervariable region, U515-E786) were prepared according to the method described by Illumina 16S metagenomic library preparation with some slight modifications (first PCR clean-up was done by using ExoSAP-IT express PCR clean up reagent, Thermo Fisher). Samples were sequenced on an Illumina MiSeq (PE 250+250) at the BioMicro Center (Massachusetts Institute of Technology, Cambridge, MA). Reads were processed using a custom pipeline where cutadapt was used for primer trimming, QIIME 1.9 (Caporaso et al., 2010) was used for demultiplexing, and DADA2 (Callahan et al., 2016)was used to infer amplicon sequence variants (ASVs). 16S copy number correction was performed with microbiome helper; sequence variants were combined with the Greengenes database v13.5(DeSantis et al., 2006) to build a new reference tree using FastTree, and assigned copynumbers using PICRUST(Langille et al., 2013). Taxonomy for the sequence variants was assigned using the RDP database(Cole et al., 2014).

### Calculation of RTF_C_

For *RTF*_*C*_ calculation, the monoculture functions measured from the second dilution regrowth was used as *F*_*i*_.*F*_*C,i*_ was calculated from *F*_*C*_*R*_*C,i*_, where *F*_*C*_ is the total community function and *R*_*C,i*_ is the relative abundance of the of taxon *i* in community *C*. This is only completely accurate when the community function of study is cell density, since different taxa may have different function to cell ratios. Thus, for community functions other than cell density,

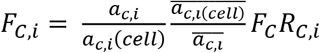

*F*_*C*_*R*_*C,i*_ can be used as an approximation for *F*_*C,i*_ as long as the adjustment factor 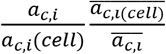 is close to 1. Under the null model, the adjustment factor becomes 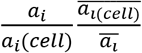, which we find to be between the ranges of 0.5-2 for the majority of our taxa in communities (Figure S8). The overall effect of this adjustment factor on *RTF*_*C*_ is further averaged when summed across different taxa in a community. Thus, for simplicity, the adjustment factor was assumed to be 1 for all taxa in communities, i.e. the protein/cell ratio and CO_2_/cell ratio among taxa in a community were equal.

Other estimations and assumptions used to calculate RTF included: i) The criteria for determining if a sample was a monoculture was that 90% of the reads in the sample belonged to one sequence variant; the functions of the community was directly used as the monoculture functions. ii) The criteria for a community to be “constitutable” was that 85% of the reads were covered by sequence variants that we had a monoculture function of. The constitutable parts of the communities were re-normalized so that the relative abundances of the sequence variants added up to 100%. iii) CO_2_ production from 0-40h were used for *RTF*_*C*_ calculations for CO_2_, since community composition measurements taken at early stationary phase were all around 40h.

### Estimation of relative CUE

Since 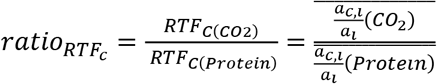, and 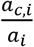 is a ratio, we can alter the units of *a*_*i*_ so that *a*_*i*(*Protien*)_ and 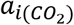 are the same units. Now *a*_*i*(*Protien*)_ is equivalent to CUE, and 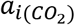 is equivalent to 1-CUE. By definition, *a*_*C,i*(*Protien*)_ is relative-to-expected CUE (denoted as R) times CUE. Thus, by seeing the whole community as the behavior as an “average taxa”, 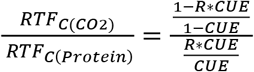. This gives

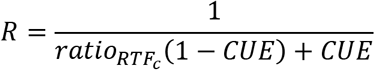

Estimation of relative CUE depending on the ratio between *RTF*_*C*(*CO2*)_ and *RTF*_*C*(*Protien*)_ was performed according to the equation above and by setting up the expected CUE in intervals of 0.01, ranging from 0-0.6.

### Curve fitting

All curve fitting were performed using the nls function in R, and the 95% confidence interval of the curve fits were performed using uncertainty propagation by first-/second-order Taylor expansion and Monte Carlo simulation including covariances using the package “propagate” (Andrej-Nikolai, 2017). A linear, log-linear, and hyperbolic least squares fit was performed for each dataset, and compared against a null model of constant value. All sum of squares calculations were type III, and performed using the function “Anova” in the R package “car”(Fox and Weisberg, 2011).

### Diversity calculations

Species richness were determined as the number of ASVs in each community, and they were compared against a rarefactioned species richness, using the R package “vegan”(Oksanen et al., 2013), and an estimated species richness, using the R package “breakaway”(Willis and Bunge, 2015). MPD and MNTDs of each community were calculated by first performing a multiple-alignment using the R package “DECIPHER”(Wright, 2016), then constructing a GTR+G+I (Generalized time-reversible with Gamma rate variation) maximum likelihood tree with the R package “phangorn”(Schliep, 2011), and calculating the actual values using the R package “picante”(Kembel et al., 2010).

---

**Figure S1.**
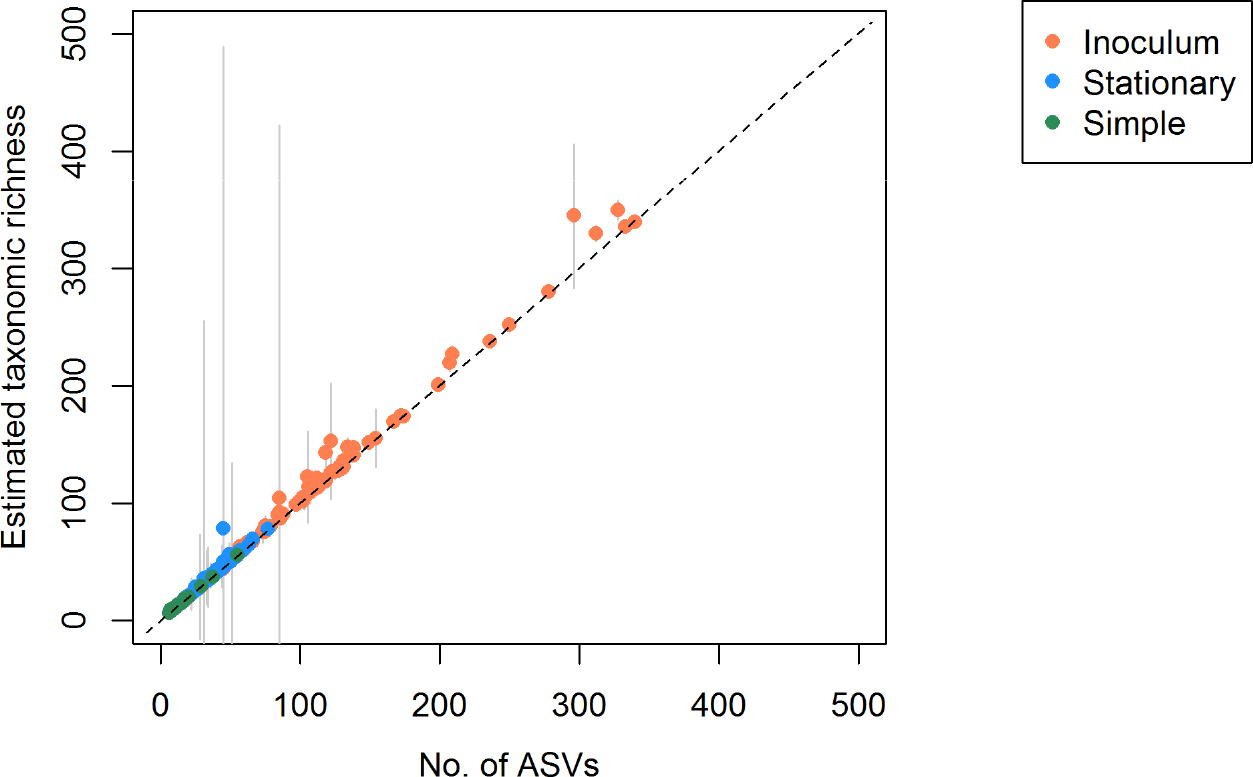
Estimated taxonomic richness are almost identical to ASv counts. Relationship between the number of ASVs counted directly from sequencing data and estimated after accounting for uncounted ASVs due to finite sequencing depth for all sequenced communities. Only communities that fit the estimation criterion (at least 6 different read frequencies) are shown. Grey bars, standard error of the estimate; broen line, estimated taxonomic richness=counted number of ASVs.

**Figure S2.**
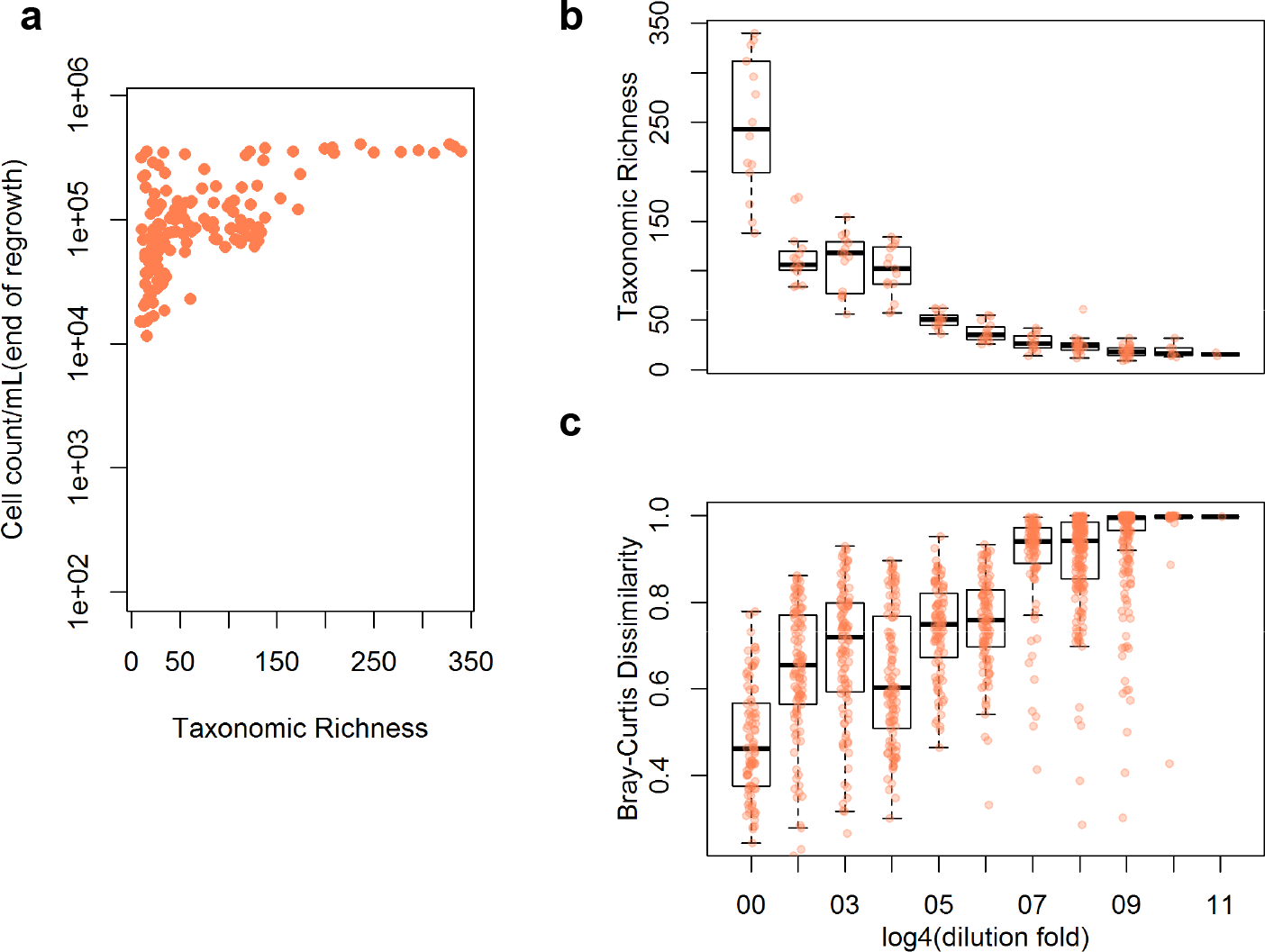
Effects of dilution-regrowth of microbial communities in pasterized seawater. (a) Taxonomic richness and cell denisty of all inoculum communities. (b) Relationship of dilution factor and taxonomic richness. (c) Bray-Curtis dissimilarity between any two communities with the same dilution factor. Each red dot represents one community; the blac line in box plots represent the median; wisers represent 95% confidence interval of the median.

**Figure S3.**
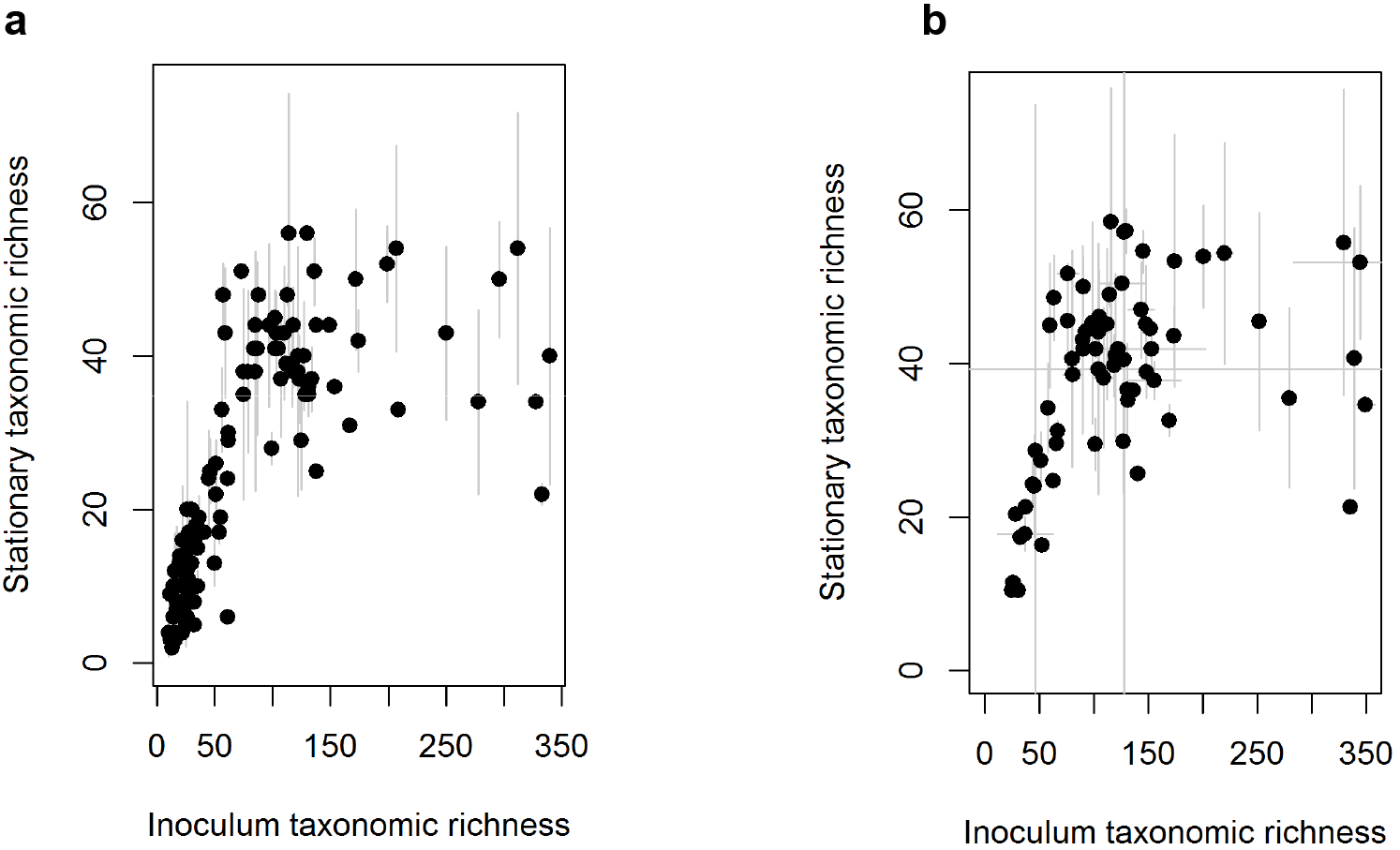
Relationship between inoculum taxonomic reichness and stationary taxonomic richness. Taxonomic richness displayed as (a) direct ASV counts and (b) estimated taxonomic richness. Each blac dot represents one inoculum community, and error bars are standard deviations over three biological replicates derived from the same inoculum community. Error bars in b) include the propagated standard error of the taxonomic richness estimates.

**Figure S4.**
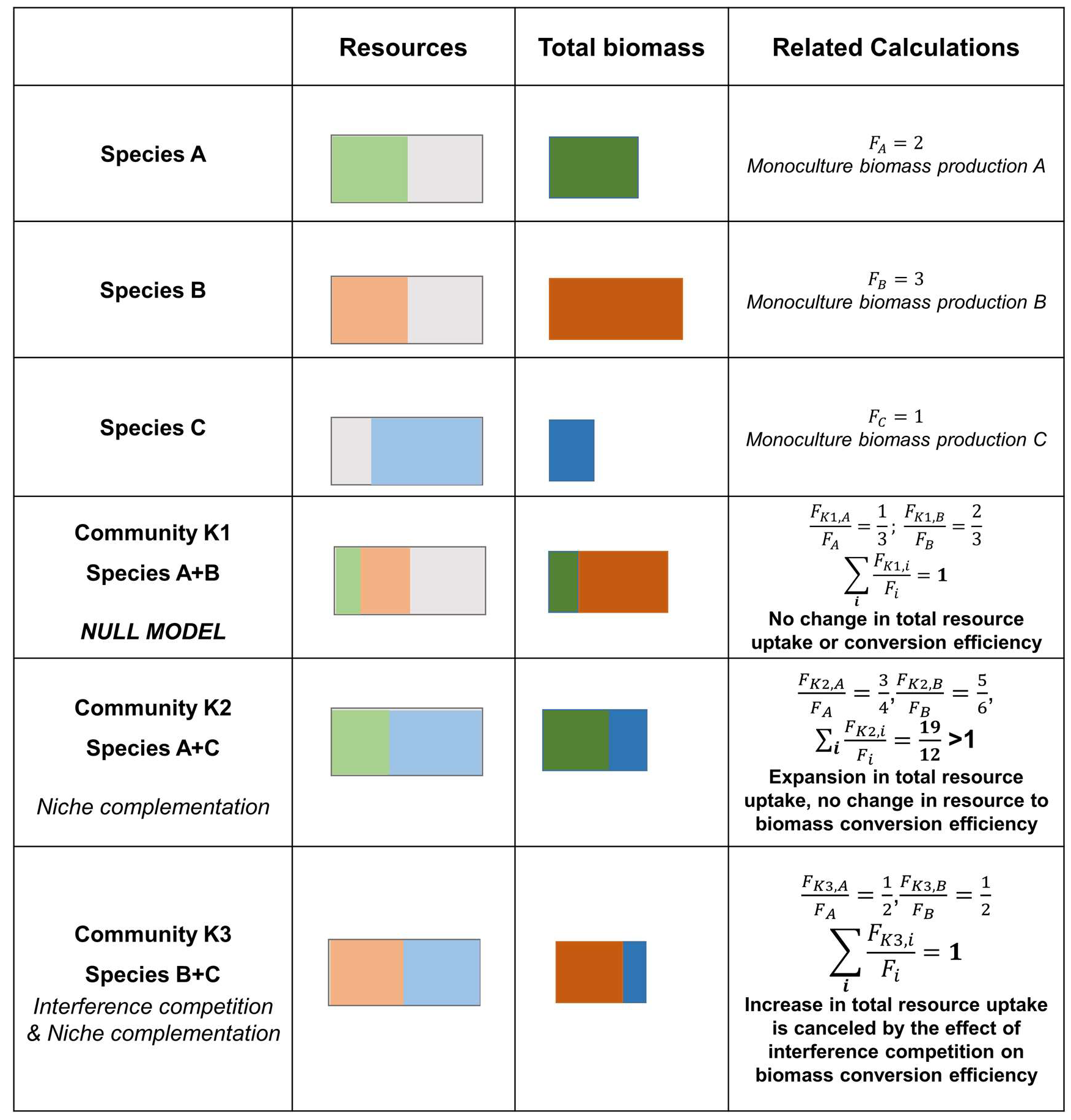
Graphical illustration of the RTF concept.

**Figure S5.**
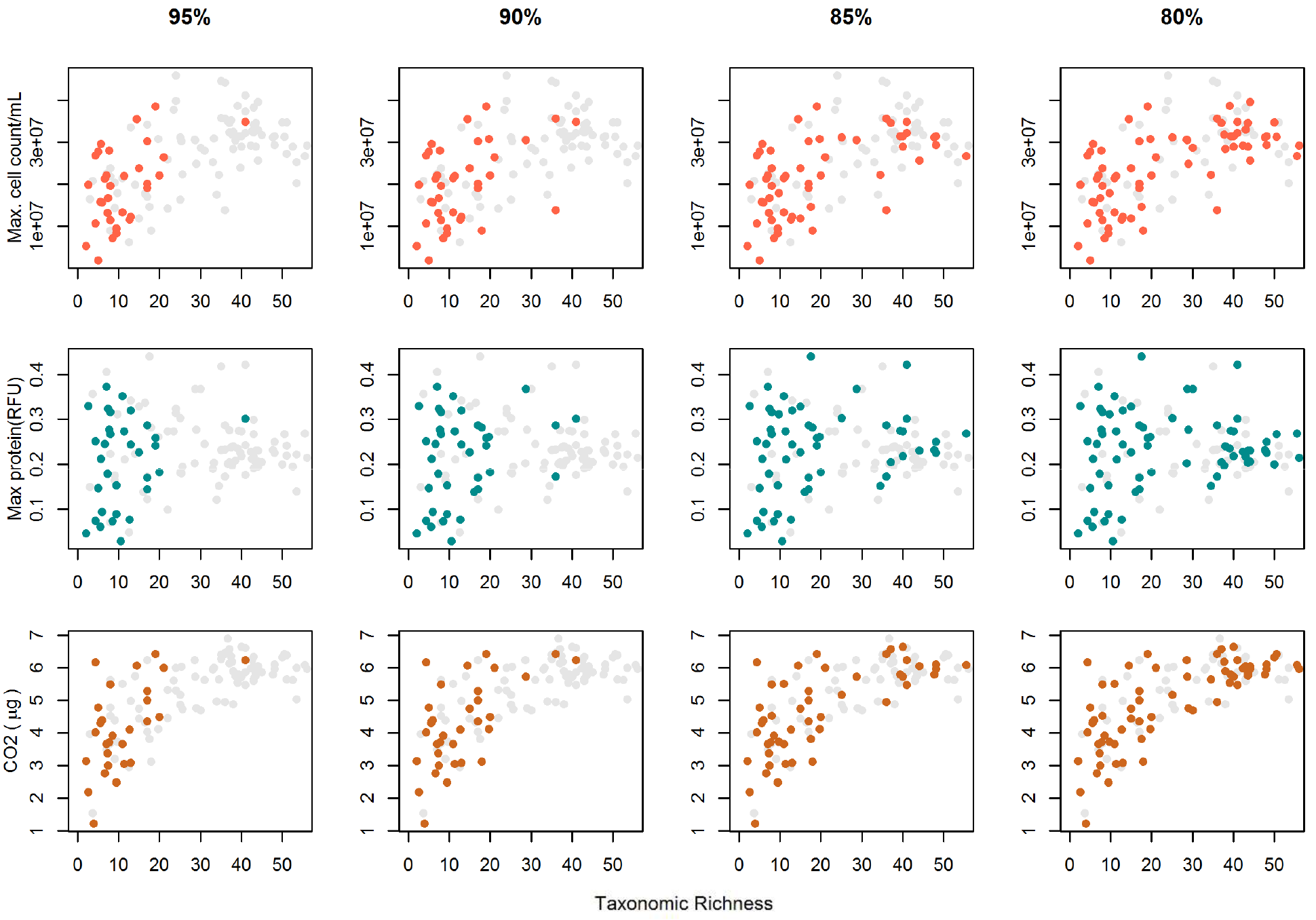
Constitutable communities under different community coverage criteria. Relaxing criteria (95%, 90%, 85%, 80% reads covered by taxa with nown monoculture productivity) for constitutable communities increase their diversity coverage. Final criteria selected for constitutable communities was 85% coverage since it was the highest standard that allowed sufficient coverage of diversity. Colored dots, communities with production and respiration measurements that are constitutable; grey dots, communities with production and respiration measurements but are not constitutable.

**Figure S6.**
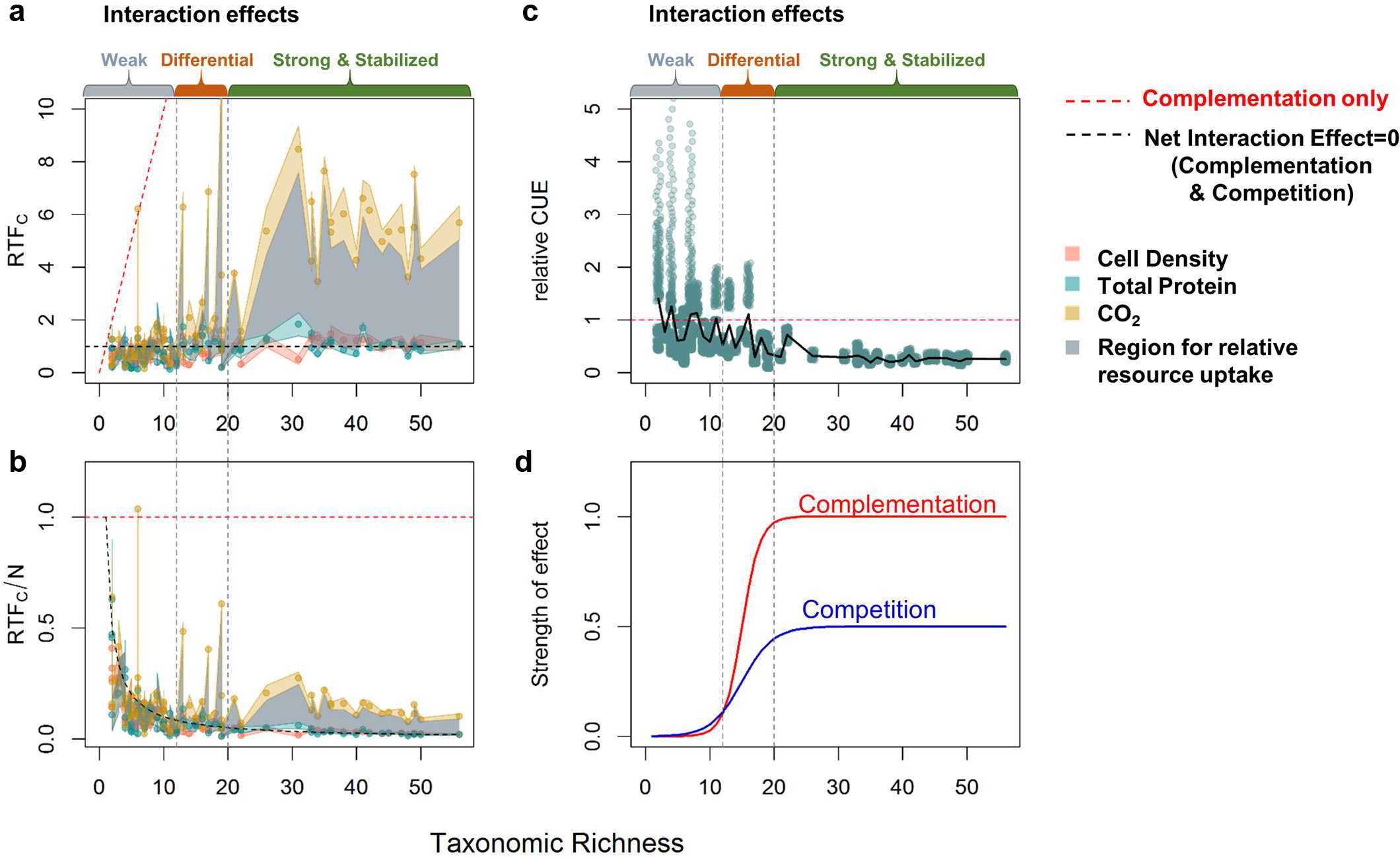
Relative community-to-monoculture functions indicate differential increases in inche complementation and competition with diversity (linear scale). Relationship between diversity and (a) relative total biomass production, respiration, and resource uptae, and (b) relative per taxa biomass production, respiration, and resource uptae (c) estimated relative carbon use efficiency (CUE). In (a), (b) each point represents one community at stationary phase (biological replicates were not averaged). In (c), each point represents a combination of one relative total function (*RTF*_*C*_) ratio with a random value of CUE drawn from 0-0.6. The blac line represents the mean of all combinations with the same taxonomic richness. Colored regions around the points indicate the standard deviation of the interaction effect, and the grey areas indicate the range where relative total and per taxa resource uptae is limited to at each taxonomic richness. (d) A hypothetical model for how the effects of niche complementation and competition on community function scales with taxonomic richness. Numbers on the y-axis are arbitrary.

**Figure S7.**
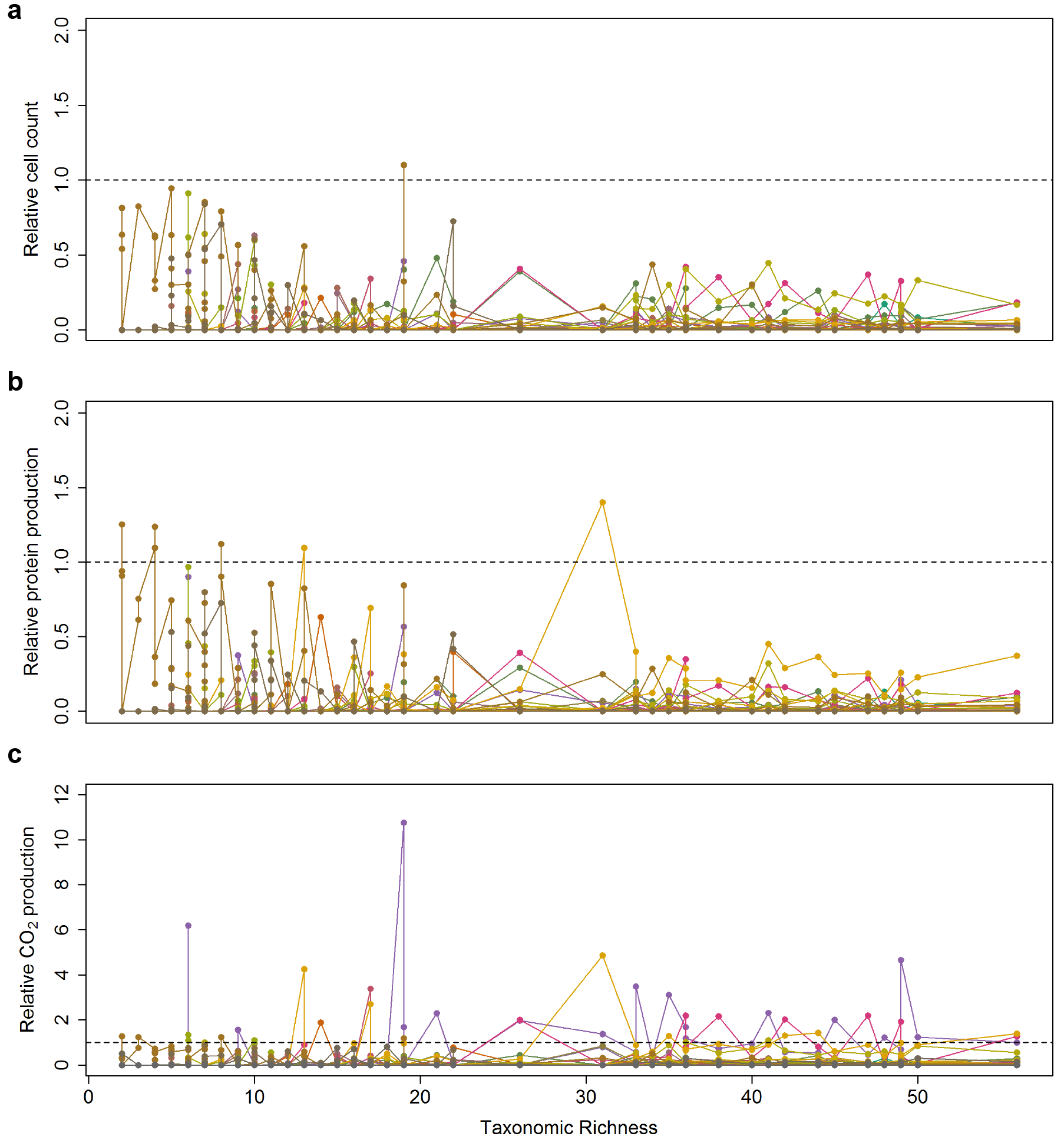
Relative function of all isolate taxa in communities. The (a) relative cell count, (b) relative protein production, and (c) relative CO_2_ production in all communities to monoculture for all the isolate taxa. Each colored line with points represents one taxa.

**Figure S8.**
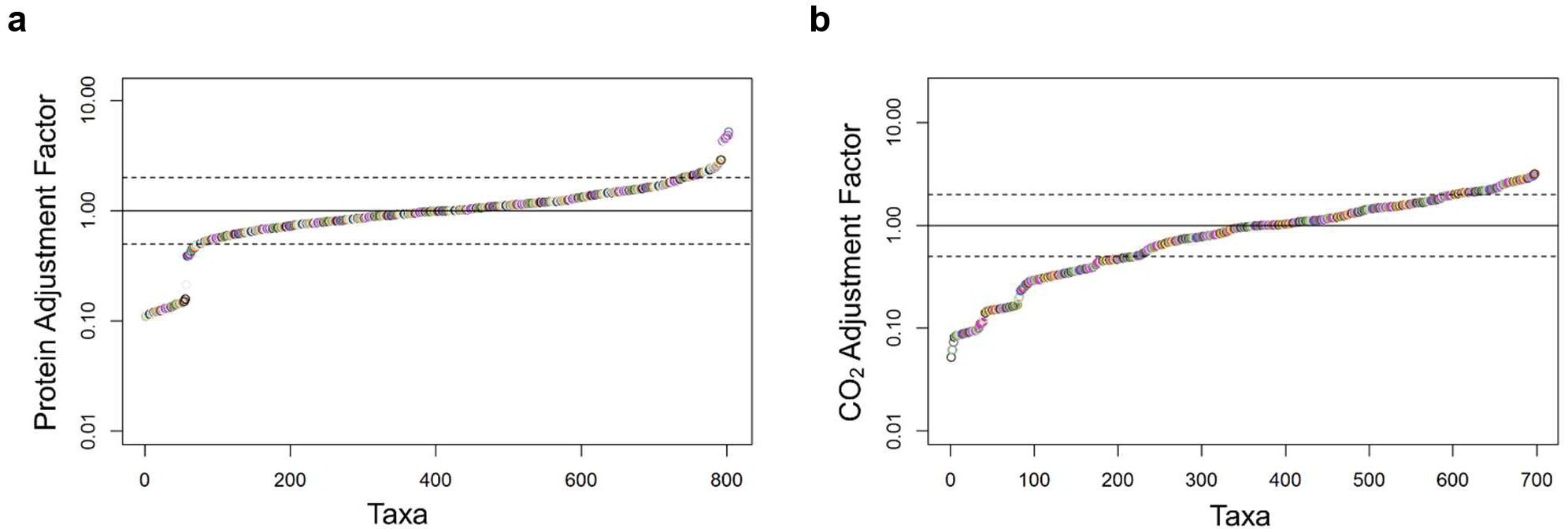
Adjustment factors for all taxa in comunitites. The adjustment factor for (a) protein measurements and (b) CO_2_ production (0-40h) for all taxa of non-zero abundance in communities. Dotted blac lines represent the range 0.5-2.

**Table S1.**
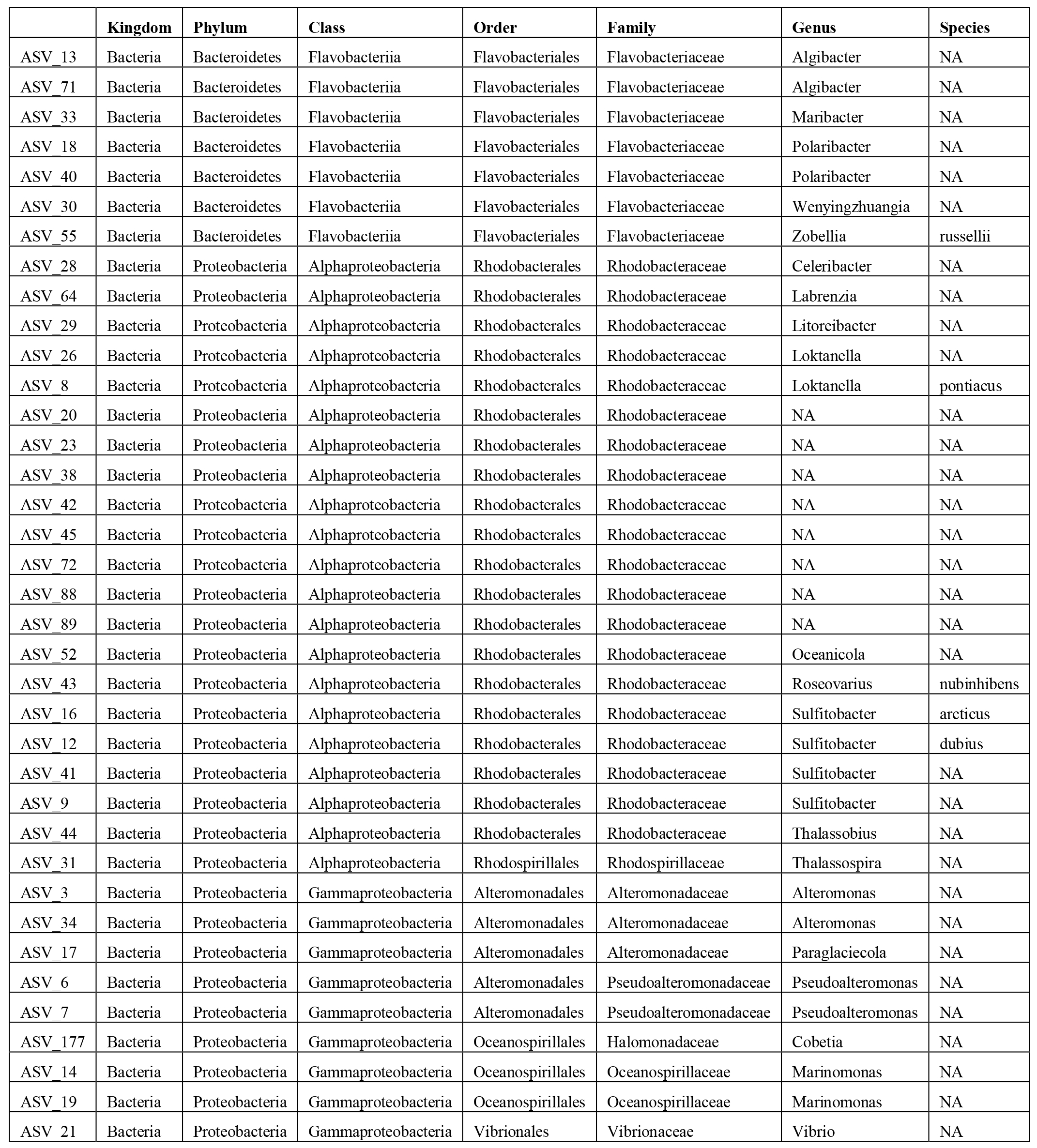
List of isolates.

**Table S2.**
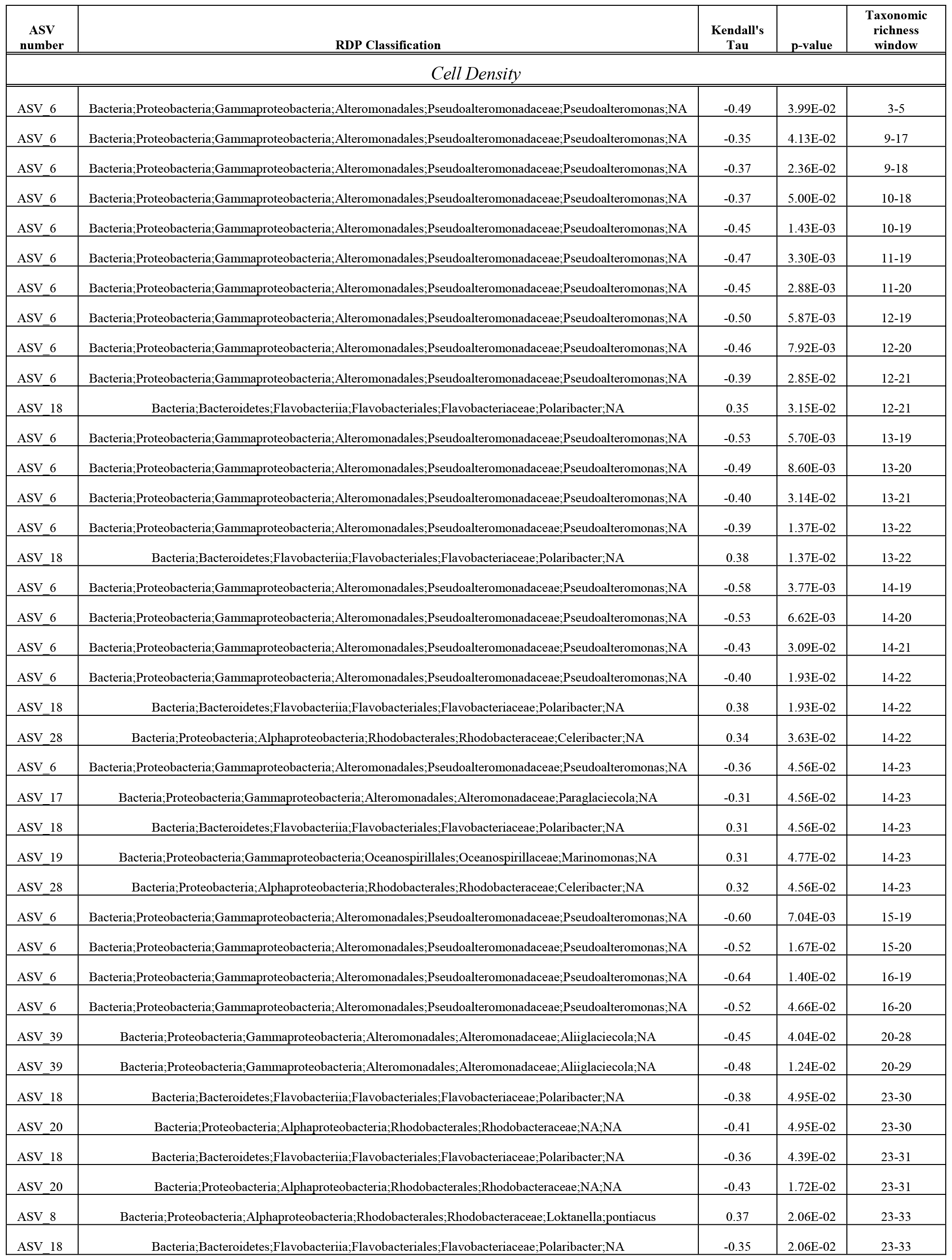
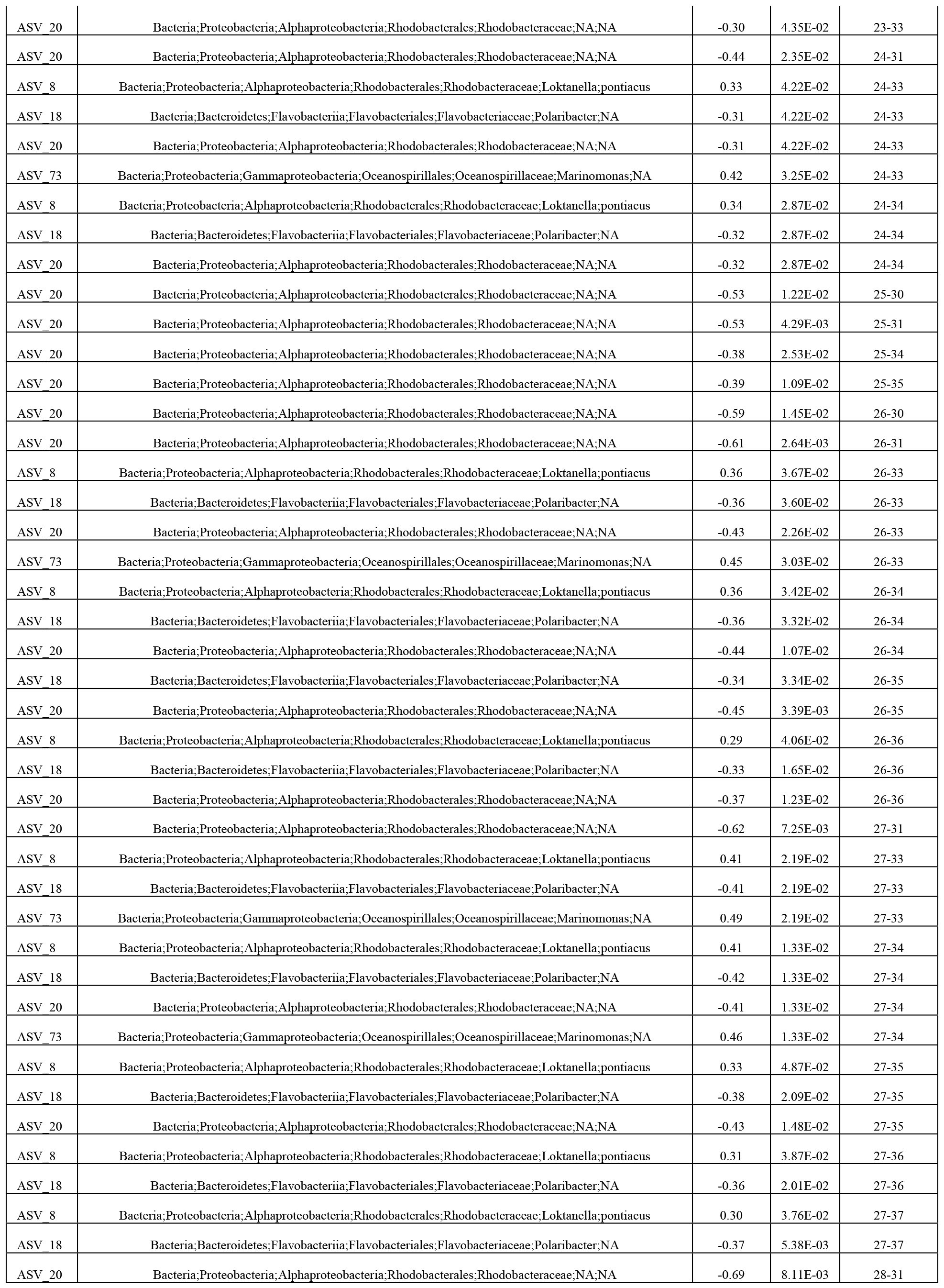
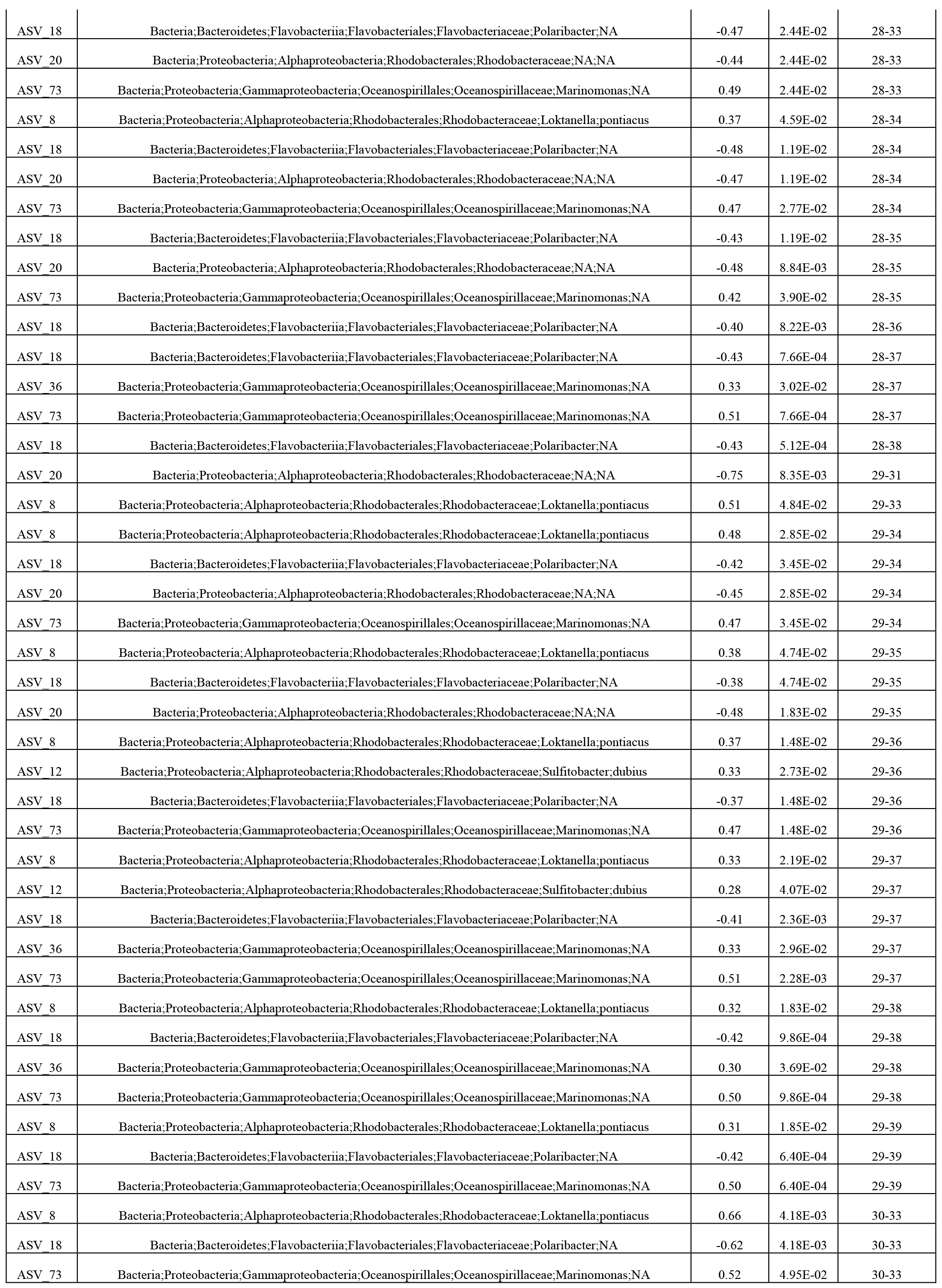
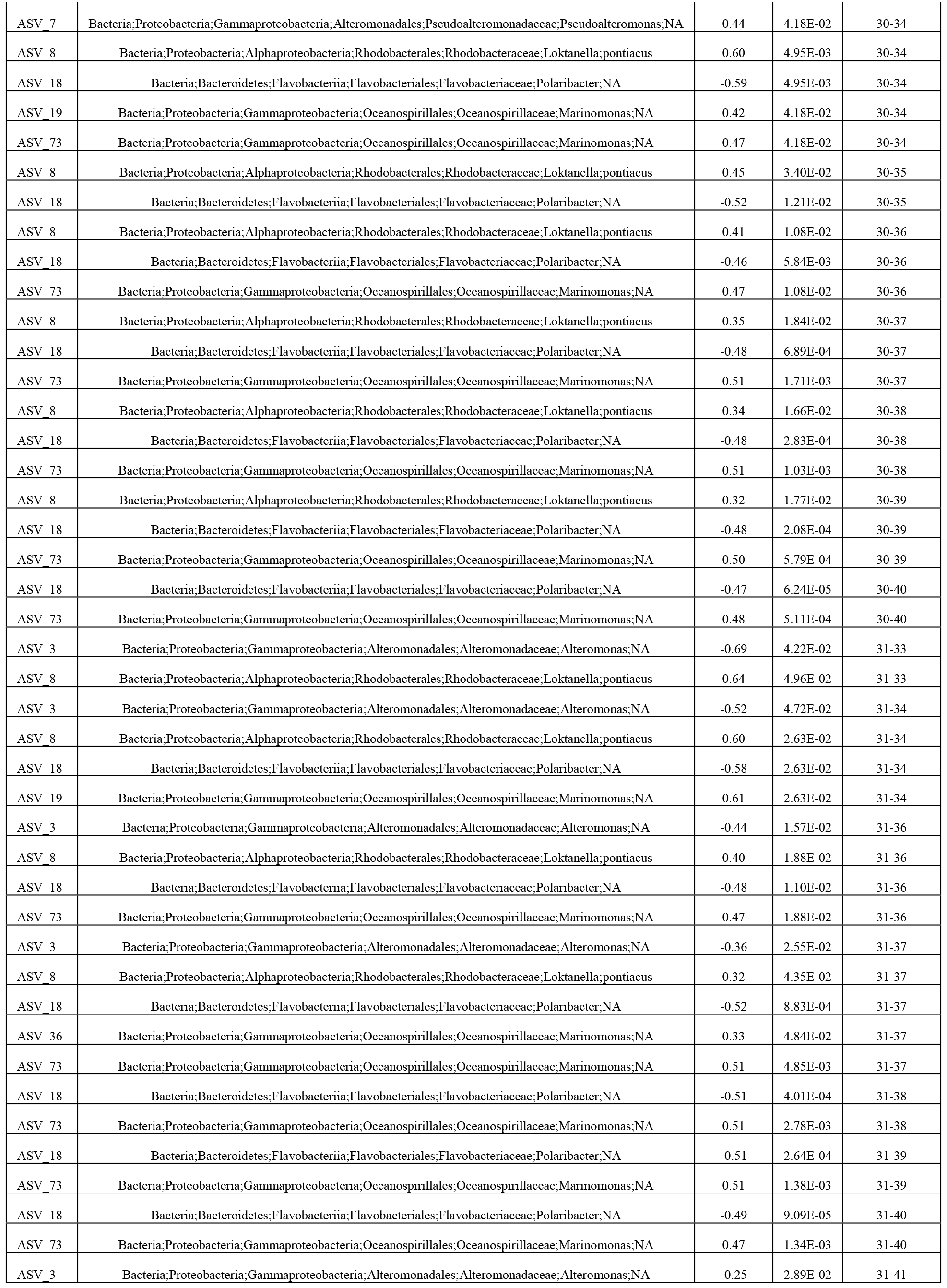
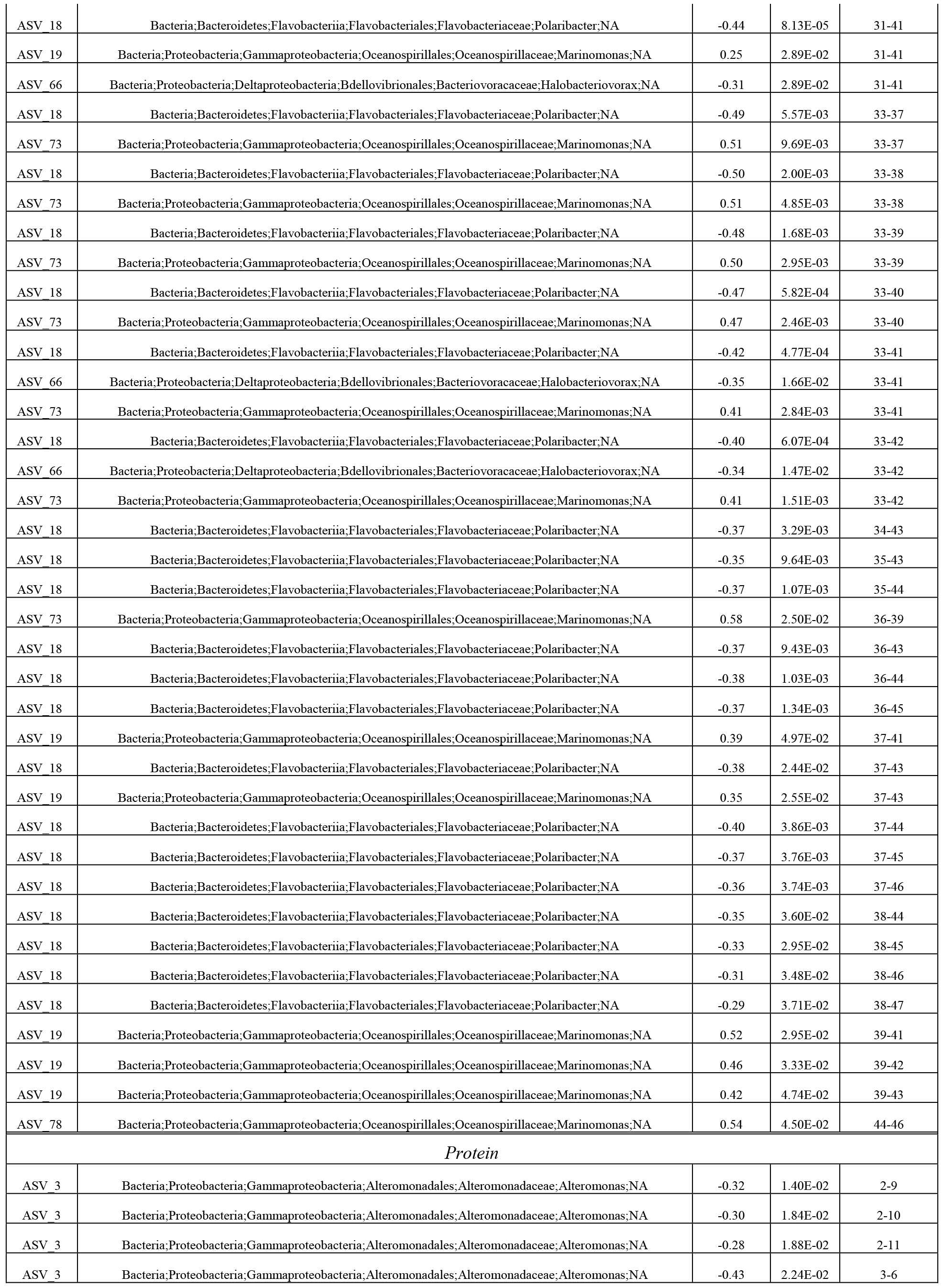
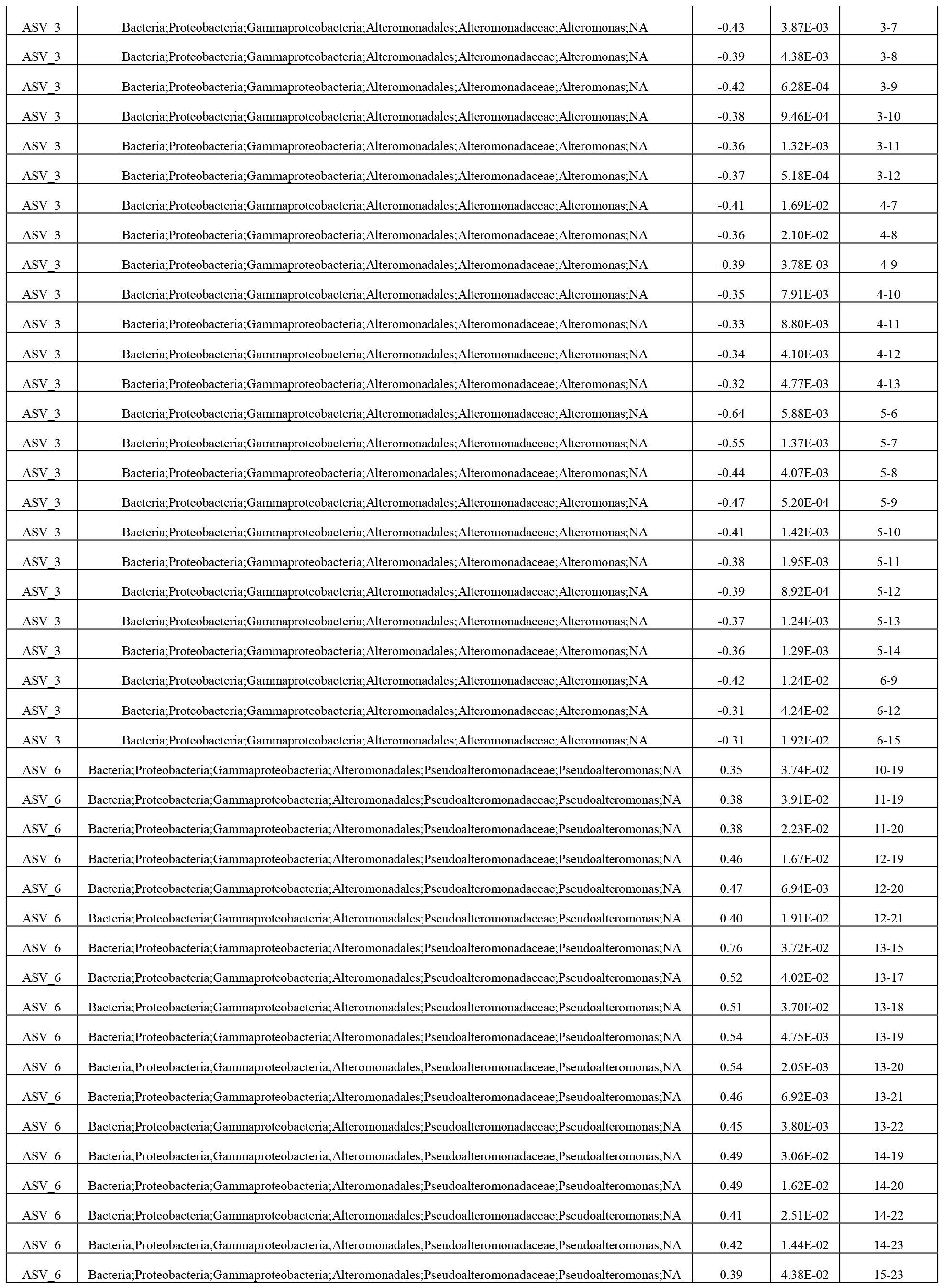
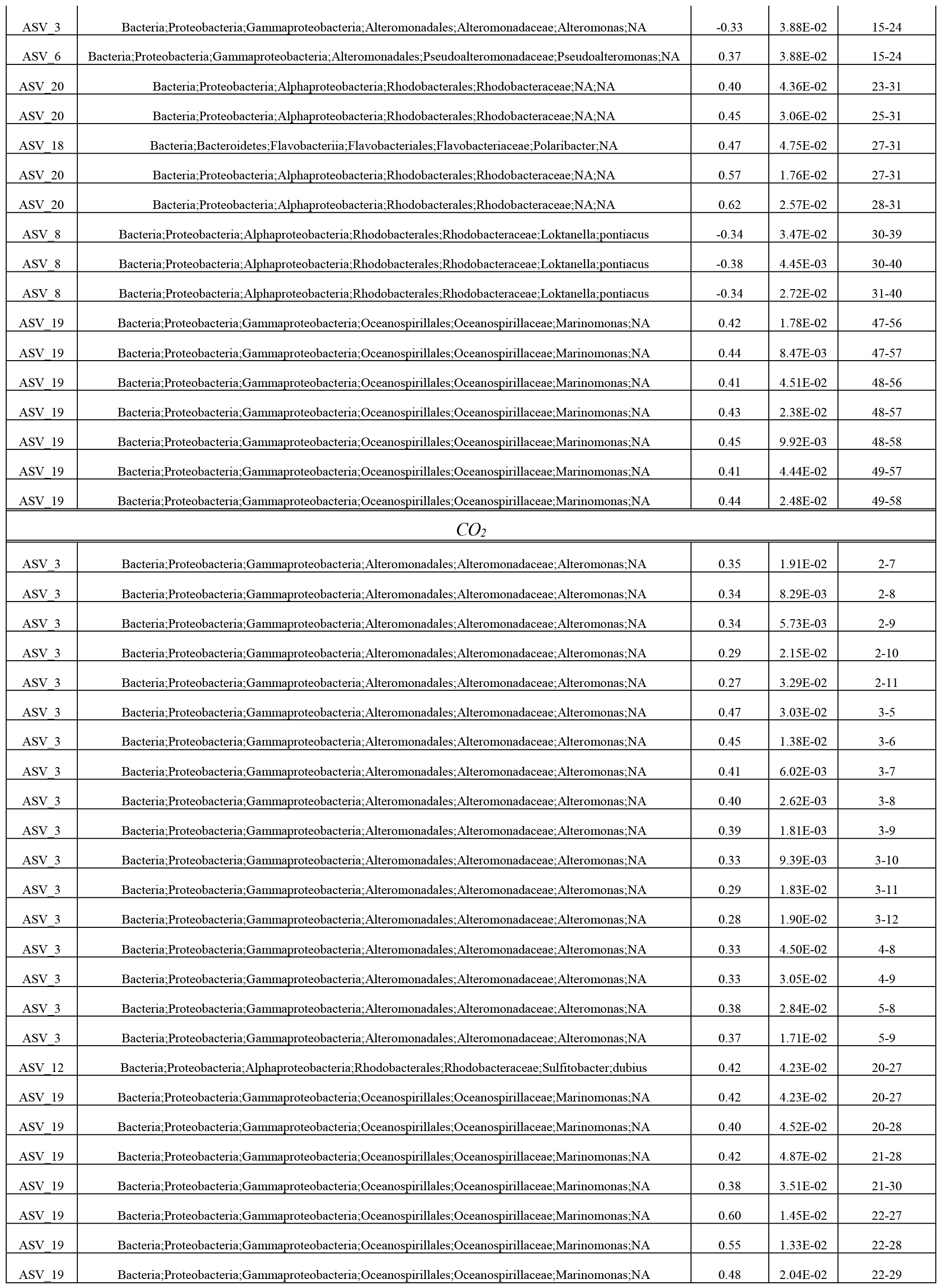
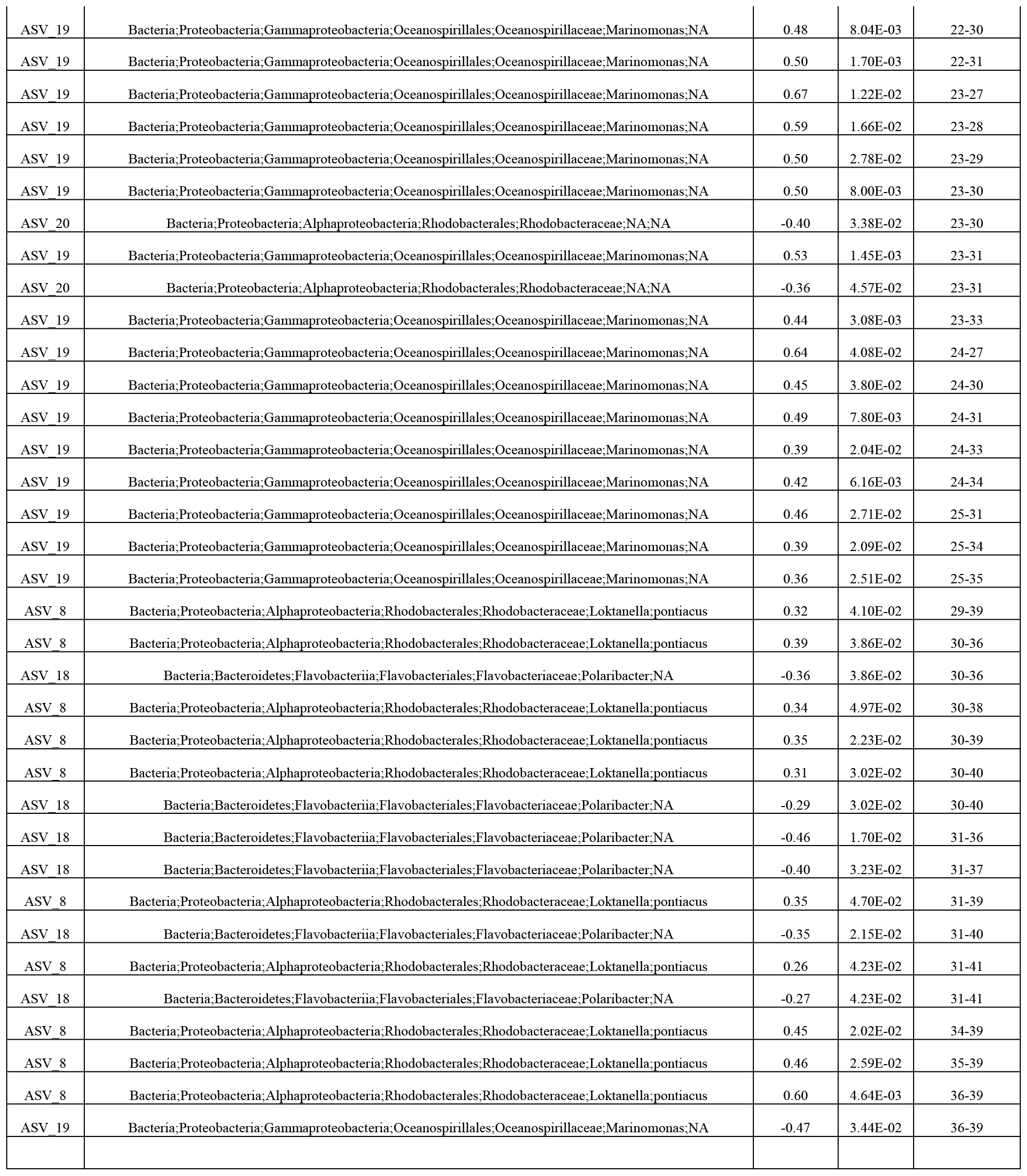
Taxa associated with community function in all taxonomic richness windows.

**Table S3.** **ANOVA of community function**

Community function in cell density
|  | Sum Sq | Df | F value | Pr(>F) |
| --- | --- | --- | --- | --- |
| log(Taxonomic richness) | 4.91E+15 | 1E+00 | 1.03E+02 | 4.1081E-20*** |
| log(Initial cell density) | 1.58E+14 | 1E+00 | 3.33E+00 | 6.9490E-02 |
| Residuals | 1.05E+16 | 2E+02 |  |  |

Community function in protein fluorescence
|  | Sum Sq | Df | F value | Pr(>F) |
| --- | --- | --- | --- | --- |
| log(Taxonomic richness) | 1.98E-02 | 1E+00 | 3.42E+00 | 6.5724E-02 |
| log(Initial cell density) | 1.23E-02 | 1E+00 | 2.12E+00 | 1.4671E-01 |
| Residuals | 1.28E+00 | 2E+02 |  |  |

Community function in CO2
|  | Sum Sq | Df | F value | Pr(>F) |
| --- | --- | --- | --- | --- |
| log(Taxonomic richness) | 1.37E+02 | 1E+00 | 2.09E+02 | 9.1034E-34 |
| log(Initial cell density) | 7.60E+00 | 1E+00 | 1.16E+01 | 7.7094E-04 |
| Residuals | 1.44E+02 | 2E+02 |  |  |

Signif. codes: \*\*\*<0.001 \*\*<0.01 \*<0.05

## References

Andrej-Nikolai Spiess (2017). propagate: Propagation of Uncertainty. R package version 1.0-5. https://CRAN.R-project.org/package=propagate

Callahan, B.J., McMurdie, P.J., Rosen, M.J., Han, A.W., Johnson, A.J.A., and Holmes, S.P. (2016). DADA2: High-resolution sample inference from Illumina amplicon data. Nat. Methods 13, 581.

Callahan, B.J., McMurdie, P.J., and Holmes, S.P. (2017). Exact sequence variants should replace operational taxonomic units in marker-gene data analysis. ISME J. 11, 2639–2643.

Caporaso, J.G., Kuczynski, J., Stombaugh, J., Bittinger, K., Bushman, F.D., Costello, E.K., Fierer, N., Peña, A.G., Goodrich, J.K., Gordon, J.I., et al. (2010). QIIME allows analysis of high-throughput community sequencing data. Nat. Methods 7, 335–336.

Cardinale, B.J., Srivastava, D.S., Emmett Duffy, J., Wright, J.P., Downing, A.L., Sankaran, M., and Jouseau, C. (2006). Effects of biodiversity on the functioning of trophic groups and ecosystems. Nature 443, 989–992.

Cermak, N., Becker, J.W., Knudsen, S.M., Chisholm, S.W., Manalis, S.R., and Polz, M.F. (2017). Direct single-cell biomass estimates for marine bacteria via Archimedes’ principle. ISME J. 11, 825–828.

Chen, F., Chang, Y., Dong, S., and Xue, C. (2016). Wenyingzhuangia fucanilytica sp. nov., a sulfated fucan utilizing bacterium isolated from shallow coastal seawater. Int. J. Syst. Evol. Microbiol. 66, 3270–3275.

Cole, J.R., Wang, Q., Fish, J.A., Chai, B., McGarrell, D.M., Sun, Y., Brown, C.T., Porras-Alfaro, A., Kuske, C.R., and Tiedje, J.M. (2014). Ribosomal Database Project: data and tools for high throughput rRNA analysis. Nucleic Acids Res. 42, D633–D642.

Cordero, O.X., Wildschutte, H., Kirkup, B., Proehl, S., Ngo, L., Hussain, F., Roux, F.L., Mincer, T., and Polz, M.F. (2012). Ecological Populations of Bacteria Act as Socially Cohesive Units of Antibiotic Production and Resistance. Science 337, 1228–1231.

C.T., de, Wit, and J.P., van den, Bergh, (1965). Competition between herbage plants. 13.

Datta, M.S., Sliwerska, E., Gore, J., Polz, M.F., and Cordero, O.X. (2016). Microbial interactions lead to rapid micro-scale successions on model marine particles. Nat. Commun. 7, ncomms11965.

Delgado-Baquerizo, M., Giaramida, L., Reich, P.B., Khachane, A.N., Hamonts, K., Edwards, C., Lawton, L.A., and Singh, B.K. (2016). Lack of functional redundancy in the relationship between microbial diversity and ecosystem functioning. J. Ecol. 104, 936–946.

DeSantis, T.Z., Hugenholtz, P., Larsen, N., Rojas, M., Brodie, E.L., Keller, K., Huber, T., Dalevi, D., Hu, P., and Andersen, G.L. (2006). Greengenes, a Chimera-Checked 16S rRNA Gene Database and Workbench Compatible with ARB. Appl. Environ. Microbiol. 72, 5069–5072.

Epstein, S. (2013). The phenomenon of microbial uncultivability. Curr. Opin. Microbiol. 16, 636–642.

Fiegna, F., Moreno-Letelier, A., Bell, T., and Barraclough, T.G. (2014). Evolution of species interactions determines microbial community productivity in new environments. ISME J.

Fletcher, H.R., Biller, P., Ross, A.B., and Adams, J.M.M. (2017). The seasonal variation of fucoidan within three species of brown macroalgae. Algal Res. 22, 79–86.

Fox, J.W. (2006). Using the price equation to partition the effects of biodiversity loss on ecosystem function. Ecology 87, 2687–2696.

Fox, J., and Weisberg, S. (2011). An R Companion to Applied Regression (SAGE Publications).

Gravel, D., Bell, T., Barbera, C., Bouvier, T., Pommier, T., Venail, P., and Mouquet, N. (2011). Experimental niche evolution alters the strength of the diversity-productivity relationship. Nature 469, 89–92.

Hehemann, J.-H., Arevalo, P., Datta, M.S., Yu, X., Corzett, C.H., Henschel, A., Preheim, S.P., Timberlake, S., Alm, E.J., and Polz, M.F. (2016). Adaptive radiation by waves of gene transfer leads to fine-scale resource partitioning in marine microbes. Nat. Commun. 7, 12860.

Jaillard, B., Richon, C., Deleporte, P., Loreau, M., and Violle, C. An a posteriori species clustering for quantifying the effects of species interactions on ecosystem functioning. Methods Ecol. Evol. n/a–n/a.

Jousset, A., Schmid, B., Scheu, S., and Eisenhauer, N. (2011). Genotypic richness and dissimilarity opposingly affect ecosystem functioning. Ecol. Lett. 14, 537–545.

Kembel, S.W., Cowan, P.D., Helmus, M.R., Cornwell, W.K., Morlon, H., Ackerly, D.D., Blomberg, S.P., and Webb, C.O. (2010). Picante: R tools for integrating phylogenies and ecology. Bioinformatics 26, 1463–1464.

Langille, M.G.I., Zaneveld, J., Caporaso, J.G., McDonald, D., Knights, D., Reyes, J.A., Clemente, J.C., Burkepile, D.E., Vega Thurber, R.L., Knight, R., et al. (2013). Predictive functional profiling of microbial communities using 16S rRNA marker gene sequences. Nat. Biotechnol. 31, 814–821.

Lipson, D.A. (2015). The complex relationship between microbial growth rate and yield and its implications for ecosystem processes. Front. Microbiol. 6.

Loreau, M., and Hector, A. (2001). Partitioning selection and complementarity in biodiversity experiments. Nature 412, 72–76.

Mas-Lladó, M., Piña-Villalonga, J.M., Brunet-Galmés, I., Nogales, B., and Bosch, R. (2014). Draft Genome Sequences of Two Isolates of the Roseobacter Group, Sulfitobacter sp. Strains 3SOLIMAR09 and 1FIGIMAR09, from Harbors of Mallorca Island (Mediterranean Sea). Genome Announc. 2.

May, R.M., and Arthur, R.H.M. (1972). Niche Overlap as a Function of Environmental Variability. Proc. Natl. Acad. Sci. 69, 1109–1113.

Maynard, D.S., Crowther, T.W., and Bradford, M.A. (2017). Competitive network determines the direction of the diversity-function relationship. Proc. Natl. Acad. Sci. 201712211.

Maynard, D.S., Crowther, T.W., and Bradford, M.A. Fungal interactions reduce carbon use efficiency. Ecol. Lett. n/a–n/a.

Oksanen, J., Blanchet, F.G., Kindt, R., Legendre, P., Minchin, P., O’Hara, R., Simpson, G., Solymos, P., Stevens, M., and Wagner, H. (2013). Vegan: Community Ecology Package. R Package Version. 2.0-10. CRAN.

Pedler, B.E., Aluwihare, L.I., and Azam, F. (2014). Single bacterial strain capable of significant contribution to carbon cycling in the surface ocean. Proc. Natl. Acad. Sci. 111, 7202–7207.

Pfeiffer, T., Schuster, S., and Bonhoeffer, S. (2001). Cooperation and Competition in the Evolution of ATP-Producing Pathways. Science 292, 504–507.

Polz, M.F., and Cordero, O.X. (2016). Bacterial evolution: Genomics of metabolic trade-offs. Nat. Microbiol. 1, 16181.

Roller, B.R.K., Stoddard, S.F., and Schmidt, T.M. (2016). Exploiting rRNA operon copy number to investigate bacterial reproductive strategies. Nat. Microbiol. 1, 16160.

Rypien, K.L., Ward, J.R., and Azam, F. (2010). Antagonistic interactions among coral-associated bacteria. Environ. Microbiol. 12, 28–39.

Schaechter, M., Maaloe, O., and Kjeldgaard, N.O. (1958). Dependency on medium and temperature of cell size and chemical composition during balanced grown of Salmonella typhimurium. J. Gen. Microbiol. 19, 592–606.

Schliep, K.P. (2011). phangorn: phylogenetic analysis in R. Bioinformatics 27, 592–593.

Shen, J., Chang, Y., Dong, S., and Chen, F. (2017). Cloning, expression and characterization of a ι-carrageenase from marine bacterium Wenyingzhuangia fucanilytica: A biocatalyst for producing ι-carrageenan oligosaccharides. J. Biotechnol. 259, 103–109.

Singh, R.P., and Reddy, C.R.K. (2014). Seaweed-microbial interactions: key functions of seaweed-associated bacteria. FEMS Microbiol. Ecol. 88, 213–230.

Takemura, A.F., Corzett, C.H., Hussain, F., Arevalo, P., Datta, M., Yu, X., Le Roux, F., and Polz, M.F. (2017). Natural resource landscapes of a marine bacterium reveal distinct fitness-determining genes across the genome. Environ. Microbiol. 19, 2422–2433.

Teeling, H., Fuchs, B.M., Becher, D., Klockow, C., Gardebrecht, A., Bennke, C.M., Kassabgy, M., Huang, S., Mann, A.J., Waldmann, J., et al. (2012). Substrate-Controlled Succession of Marine Bacterioplankton Populations Induced by a Phytoplankton Bloom. Science 336, 608–611.

Willis, A., and Bunge, J. (2015). Estimating diversity via frequency ratios. Biometrics 71, 1042–1049.

Wright, E.S. (2016). Using DECIPHER v2.0 to Analyze Big Biological Sequence Data in R. 8, 8.

Xing, P., Hahnke, R.L., Unfried, F., Markert, S., Huang, S., Barbeyron, T., Harder, J., Becher, D., Schweder, T., Glöckner, F.O., et al. (2015). Niches of two polysaccharide-degrading Polaribacter isolates from the North Sea during a spring diatom bloom. ISME J. 9, 1410–1422.

